# Covalent Degrader of the Oncogenic Transcription Factor β-Catenin

**DOI:** 10.1101/2023.10.31.565018

**Authors:** Flor A. Gowans, Nafsika Forte, Justin Hatcher, Oscar W. Huang, Yangzhi Wang, Belen E. Altamirano Poblano, Ingrid E. Wertz, Daniel K. Nomura

**Author notes:** co-first authors that contributed equally to this work (Flor A. Gowans, Nafsika Forte).

## Abstract

β-catenin (CTNNB1) is an oncogenic transcription factor that is important in cell-cell adhesion and transcription of cell proliferation and survival genes that drives the pathogenesis of many different types of cancers. However, direct pharmacological targeting of CTNNB1 has remained challenging deeming this transcription factor as “undruggable.” Here, we have performed a screen with a library of cysteine-reactive covalent ligands to identify a monovalent degrader EN83 that depletes CTNNB1 in a ubiquitin-proteasome-dependent manner. We show that EN83 directly and covalently targets CTNNB1 through targeting four distinct cysteines within the armadillo repeat domain—C439, C466, C520, and C619—leading to a destabilization of CTNNB1. Using covalent chemoproteomic approaches, we show that EN83 directly engages CTNNB1 in cells with a moderate degree of selectivity. We further demonstrate that direct covalent targeting of three of these four cysteines--C466, C520, and C619--in cells contributes to CTNNB1 degradation in cells. We also demonstrate that EN83 can be further optimized to yield more potent CTNNB1 binders and degraders. Our results show that chemoproteomic approaches can be used to covalently target and degrade challenging transcription factors like CTNNB1 through a destabilization-mediated degradation.

## Introduction

β-catenin (CTNNB1) is an oncogenic transcription factor that drives the pathogenesis of many different types of human cancers, including liver, lung, colorectal, breast, and ovarian cancers ^1–3^. CTNNB1 regulates cell-cell adhesion as part of a larger protein complex with E-cadherin and α-catenin. CTNNB1 is also a transcription factor that is activated by upstream Wnt signaling pathways that regulates genes involved in cell proliferation, survival, epithelial-to-mesenchymal transition, migration, and metastasis^1–3^. CTNNB1 is highly regulated by the ubiquitin-proteasome system, wherein the E3 ubiquitin ligase β-TrCP1 recognizes N-terminus of CTNNB1 upon phosphorylation by GSK3α and GSK3β^1–3^. Mutations in CTNNB1 are commonly found in a variety of cancers that are often located in the N-terminal segment that is recognized by E3 ligases preventing ubiquitination and degradation of CTNNB1, thereby enhancing CTNNB1 oncogenic transcriptional activity^1–3^. Mutations have also been found in the Wnt pathway, including in APC and axin that act to enhance CTNNB1 levels and activity^1–3^.

Many efforts have been made to target the Wnt pathway for cancer therapy. PKF115-584 and CGP04090 disrupt interactions between CTNNB1 and TCF ^4,5^. ICG-001 disrupts binding between CTNNB1 and CBP ^6^. Tankyrase inhibitors, such as IWR-1 and XAV939, stabilize Axin and the CK1 activator Pryivinium to enhance the activity of the destruction complex have been discovered to enhance CTNNB1 degradation ^7,8^. Direct CTNNB1 binding degraders such as methyl 3-{[(4-methylphenyl)sulfonyl]amino}benzoate) or MSAB that attenuate colorectal tumor growth have also been discovered. However, given the rapid turnover rate of CTNNB1, more durable pharmacological strategies are needed to directly and fully target, degrade, and inhibit the activity of CTNNB1 for cancer therapy ^9^. Direct targeting of oncogenic transcription factors like CTNNB1 has also remained challenging due to its structural flexibility, intrinsic disorder, and lack of ligandable hotspots^2^.

There has been a recent surge in covalent drug discovery because of the ability of covalent drugs to access classically undruggable targets as well as shallower binding pockets through a combination of reactivity and affinity-driven mechanisms ^10^. With advances in covalent chemoproteomic strategies such as activity-based protein profiling (ABPP), the overall target engagement and proteome-wide selectivity of these compounds can also be assessed and coupled with medicinal chemistry efforts to not only optimize potency, but also specificity to eventually reduce off-target toxicological liabilities ^11–14^. Previously, covalent chemoproteomic approaches were also used to covalently target an intrinsically disordered cysteine within MYC to destabilize and degrade MYC leading to antitumorigenic effects ^15^, indicating that covalent ligands may uniquely enable targeting of other oncogenic transcription factors such as CTNNB1.

In this study, we have screened a cysteine-reactive covalent ligand library to discover a covalent monovalent degrader of CTNNB1 that eliminates CTNNB1 in a ubiquitin-proteasome dependent manner through targeting several cysteines on CTNNB1.

## Results

### Discovery of a Covalent Degrader of CTNNB1

To identify a covalent degrader of CTNNB1, we screened a library of 2100 cysteine-reactive covalent ligands consisting of acrylamides and chloroacetamides in HEK293 cells expressing CTNNB1 with a N-terminal HiBiT tag incorporated into the endogenous loci of CTNNB1 **(Figure 1a, Table S1)**. We identified 17 hits that lowered CTNNB1 levels. We subsequently eliminated hits that had appeared in previous screens conducted in our lab and counterscreened any resulting hits to identify compounds that showed attenuation in CTNNB1 loss upon pre-incubation with either a proteasome inhibitor bortezomib or the Cullin E3 ligase NEDDylation inhibitor MLN4924. We identified EN83 as the only hit that showed dose-responsive CTNNB1 loss that was significantly attenuated by proteasome or NEDDylation inhibition **(Figure 1b-1d)**. We further confirmed proteasome-mediated loss of CTNNB1 by Western blotting in HEK293T cells **(Figure 1e-1f)**. CTNNB1 transcriptional activity was significantly inhibited in a dose-dependent manner by EN83, demonstrating functional inhibition of the WNT pathway in cells **(Figure 1g)**. We observe higher potency against CTNNB1 transcriptional activity in cells compared to CTNNB1 degradation, suggesting that EN83 may be engaging CTNNB1 at lower concentrations to inhibit CTNNB1 activity while degradation of CTNNB1 requires higher engagement or concentrations. While CTNNB1 mRNA levels were unchanged, EN83 treatment significantly reduced the levels of CTNNB1 target gene MYC **(Figure S1)**. We also demonstrated loss of CTNNB1 in colorectal cancer cells that are driven by CTNNB1, including HT29, COLO, and SW480 cancer cell lines **(Figure 1h)**. We note that while we observed CTNNB1 loss in the HiBiT-tagged CTNNB1-expressing HEK293T cells at 24 h, we only observed CTNNB1 loss in HT29, COLO, and SW480 cells at earlier acute timepoints of 2 and 4 h, but CTNNB1 levels had recovered by 24 h in these cancer cell lines. This may be due to more rapid turn-over rates of CTNNB1 in more cancer-relevant cell lines.

**Figure 1.**
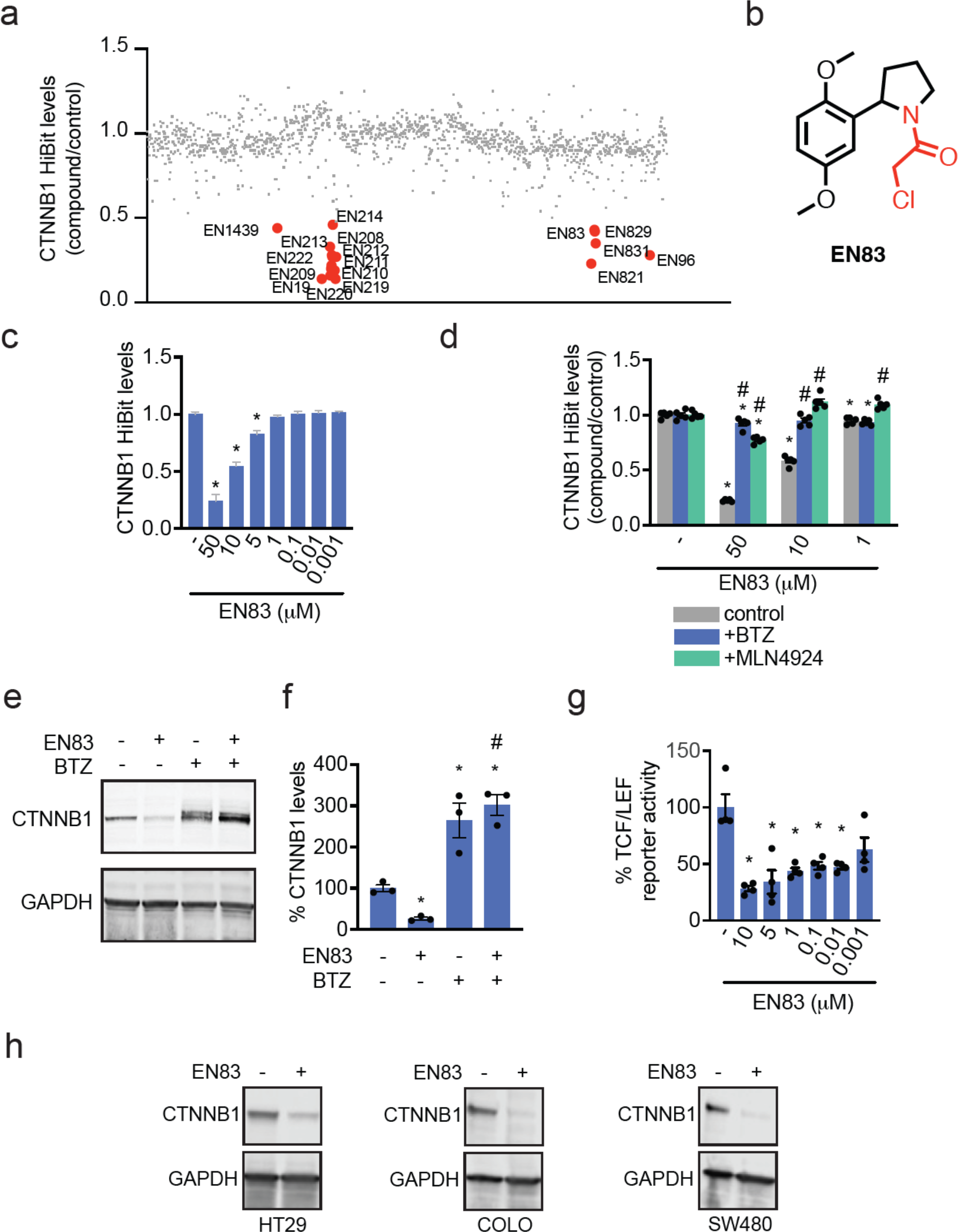
Covalent ligand screening to identify covalent CTNNB1 degraders. **(a)** Covalent ligand screen to identify compounds that lower CTNNB1 levels in cells. HEK293 cells expressing a N-terminal HiBiT tagged CTNNB1 in the endogenous CTNNB1 locus were treated with DMSO vehicle or a cysteine-reactive covalent ligand (50 μM) for 24 h and HiBiT-CTNNB1 levels were quantified. Data is shown as ratio of compound treatment/DMSO. Shown in red are compounds where there was >50 % reduction in HiBiT-CTNNB1 levels. **(b)** Structure of main hit EN83 that showed reproducible lowering of HiBiT-CTNNB1 levels where the loss of CTNNB1 was attenuated by a proteasome inhibitor. **(c)** Dose-responsive lowering of HiBiT-CTNNB1 levels with EN83 treatment. HiBiT-CTNNB1 HEK293 cells were treated with DMSO vehicle or EN83 for 24 h. **(d)** Attenuation of EN83-mediated lowering of HiBiT-CTNNB1 levels upon proteasome and NEDDylation inhibitor pre-treatment. HiBiT-CTNNB1 HEK293 cells were pre-treated with DMSO vehicle, proteasome inhibitor bortezomib (1 μM), or NEDDylation inhibitor MLN4924 (1 μM) for 1 h prior to treatment of cells with DMSO vehicle or EN83 for 24 h. **(e)** EN83-mediated lowering of CTNNB1 protein levels is attenuated by proteasome inhibitor. Conditions of treatment were as described in **(d)** but CTNNB1 and loading control GAPDH levels were assessed by Western blotting. **(f)** Quantification of experiment from **(e). (g)** TCF/LEF luciferase reporter activity of WNT/CTNNB1 transcriptional activity in HEK293 cells. HEK293 cells were transfected with the TCF/LEF luciferase reporter construct and then treated with DMSO vehicle or EN83 for 24 h after which luciferase activity was quantified. **(h)** CTNNB1 levels in colorectal cancer cells. HT29, COLO, and SW480 colorectal cancer cells were treated with DMSO vehicle or EN83 (50 μM) for 2 h. Cells that had lost adherence but were still viable were collected and CTNNB1 and loading control GAPDH levels were assessed by Western blotting. Blots and data in **(c-h)** are from n=3-6 biologically independent replicates/group, where blots are representative from the replicates. Bar graphs in **(c, d, f, g)** show average ± sem values and bar graphs in **(d, f, g)** also show individual replicate values. Significance **(c, d, f, g)** shown as *p<0.05 compared to DMSO-treated controls and #p<0.05 compared to EN83 treatment alone for each concentration group.

### Characterization of EN83 as a Direct CTNNB1 Binder

We next sought to determine whether EN83 directly and covalently bound to CTNNB1. We found that EN83 dose-responsively displaced cysteine-reactive fluorescent probe labeling of pure human CTNNB1 protein by gel-based ABPP **(Figure 2a)**. We further performed tandem mass spectrometry analysis (MS/MS) on tryptic digests from CTNNB1 pure protein incubated with EN83 and found four cysteines that were modified by EN83— C439, C466, C520, C619 **(Figure 2b)**. Interestingly, each of these cysteines reside in armadillo repeat motifs within the CTNNB1 structure suggesting that these ligands may be binding across several structurally similar domains. To further confirm direct covalent binding of EN83 to CTNNB1, we synthesized an alkyne-functionalized analog of EN83, NF686 and showed direct covalent dose-responsive probe labeling of pure CTNNB1 protein by NF686 **(Figure 2c-2d)**. We further demonstrated that the covalency is necessary since a non-reactive analog of EN83, NF602, does not show binding to CTNNB1 and does not alter CTNNB1 levels by HiBiT or Western blotting detection **(Figure 2f-2h)**. Our results thus compelling demonstrated that EN83 directly binds to CTNNB1.

**Figure 2.**
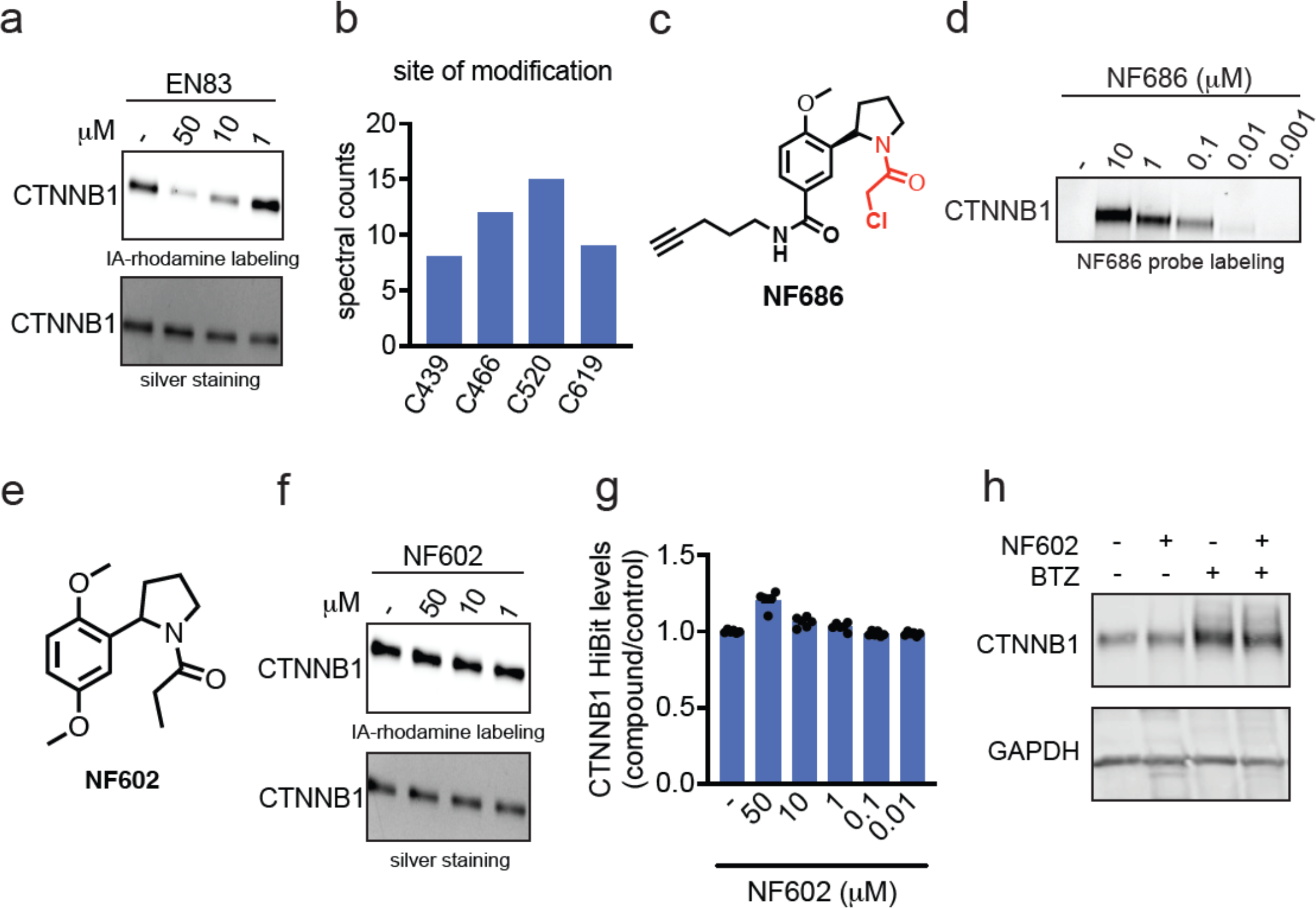
EN83 directly engages CTNNB1. **(a)** Gel-based ABPP analysis of EN83. Pure human CTNNB1 protein was pre-incubated with DMSO vehicle or EN83 for 30 min prior to labeling with a rhodamine functionalized cysteine-reactive iodoacetamide probe (IA-rhodamine) (100 nM) for 60 min after which proteins were separated on SDS/PAGE and IA-rhodamine labeling was assessed by in-gel fluorescence and protein loading was assessed by silver staining. **(b)** MS/MS analysis of EN83 site of modification on CTNNB1. Pure CTNNB1 protein was incubated with EN83 (50 μM) for 30 min after which the protein was digested with trypsin and analyzed by LC-MS/MS to look for EN83-modified cysteines. Annotated are spectral counts for four cysteines on CTNNB1 that had EN83 modifications. **(c)** Structure of alkyne-functionalized analog of EN83, NF686. **(d)** NF686 direct covalent labeling of CTNNB1. Pure human CTNNB1 protein was labeled with DMSO vehicle or NF686 for 1 h. Probe-labeled proteins were subjected to copper-catalyzed azide-alkyne cycloaddition (CuAAC) with rhodamine-azide after which proteins were separated by SDS/PAGE and probe labeling was assessed by in-gel fluorescence. **(e)** Structure of non-reactive analog of EN83, NF602. **(f)** Gel-based ABPP of NF602 performed as described in **(a). (g)** CTNNB1 levels with NF602 treatment. HiBiT-CTNNB1 HEK293 cells were treated with DMSO vehicle or NF602 for 24 h. **(h)** CTNNB1 protein levels with NF602 treatment. CTNNB1 HEK293 cells were treated with DMSO vehicle or NF602 for 24 h and CTNNB1 and loading control GAPDH levels were assessed by Western blotting. Data in **(a, b, d, f, g, h)** are from n=3-6 biologically independent replicates/group. Gels or blots shown in **(a, d, f, h)** are representative of the replicates. Data shown in **(g)** is shown as average ± sem values with individual replicates shown.

We next wanted to confirm that EN83 engages CTNNB1 in cells. We first used the alkyne probe NF686 in cells to demonstrate that CTNNB1 could be enriched from cells with the probe without enriching unrelated targets such as GAPDH **(Figure 3a)**. We next performed a mass spectrometry-based ABPP or isodesthiobiotin-ABPP (isoDTB-ABPP) to map overall proteome-wide cysteine-reactivity of EN83 in cells using previously established methods ^12,16–18^. While EN83 is only moderately selective, we did observe that EN83 significantly engaged C619 on CTNNB1 alongside 139 other targets significantly engaged by over 50 % among 8251 cysteines detected and quantified across three biological replicates **(Figure 3b, Table S2)**. Among the off-targets of EN83, none of the targets were within the Wnt signaling pathway, including AXIN1, BTRC, or GSK3A or GSK3B. Cellular thermal shift assay from EN83 cellular treatment showed significant thermal destabilization of CTNNB1 in cells, suggesting that the degradation of CTNNB1 may be caused by direct covalent targeting of CTNNB1 leading to destabilization of CTNNB1 folding and subsequent ubiquitination and degradation.

**Figure 3.**
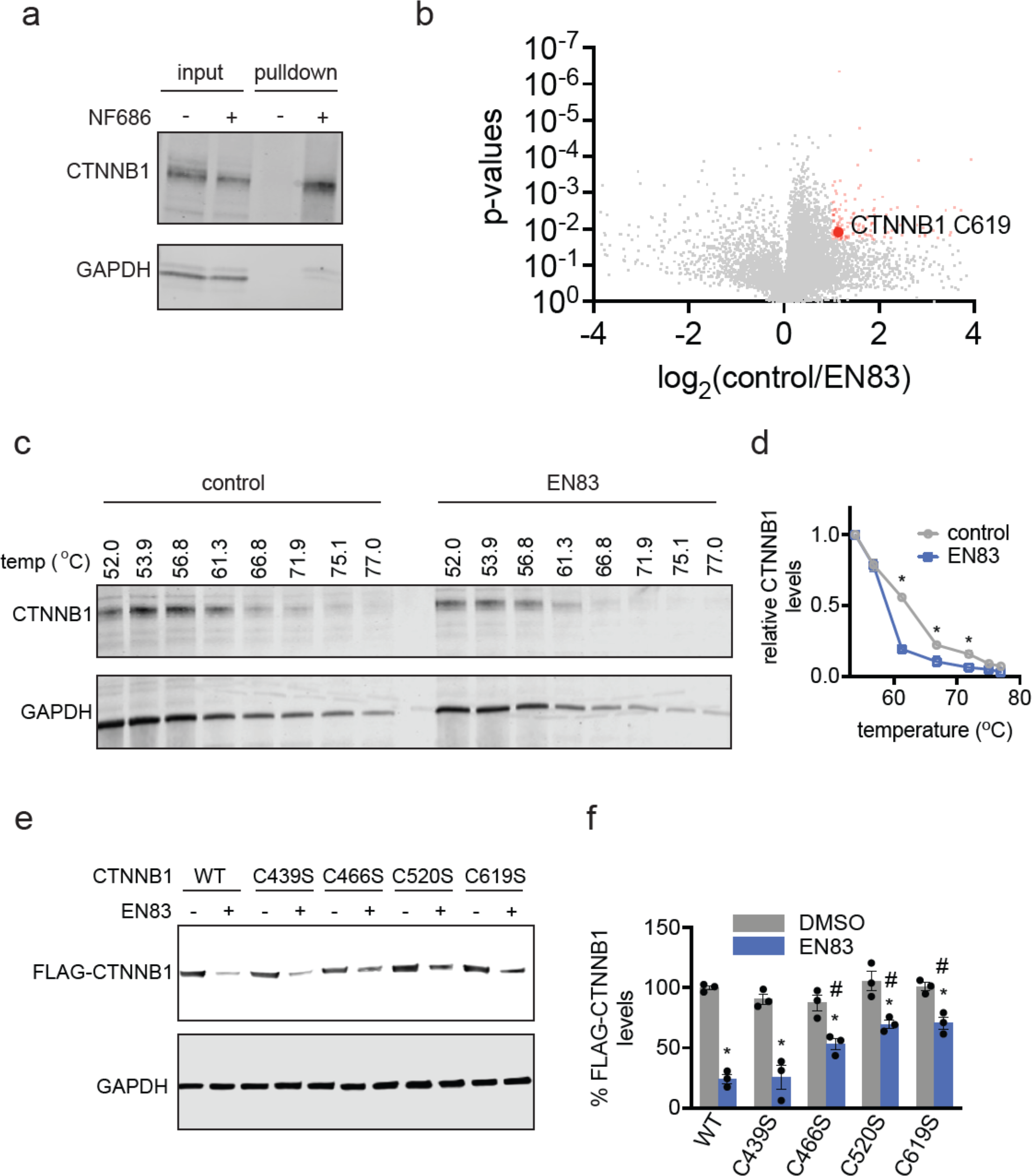
Characterization of EN83 in cells. **(a)** NF686 enrichment of CTNNB1 from cells. HiBiT-CTNNB1 HEK293 cells were treated with DMSO vehicle or NF686 (50 μM) for 24 h, after which lysates were subjected to CuAAC with biotin-azide, and probe-labeled proteins were avidin-enriched, eluted and CTNNB1 and unrelated protein GAPDH input and pulldown levels were detected by Western blotting. **(b)** Cysteine chemoproteomic profiling of EN83 by isoDTB-ABPP. HiBiT-CTNNB1 HEK293 cells were treated with DMSO vehicle or EN83 (10 μM) for 2 h. Lysates were then labeled with an alkyne-functionalized iodoacetamide probe (IA-alkyne) (200 μM) for 1 h, followed by appendage of isotopically light or heavy desthiobiotin-azide handles by CuAAC, after which probe-modified proteins were avidin-enriched, tryptically digested, and probe-modified peptides were eluted and analyzed by LC-MS/MS and control versus treated or light versus heavy probe-modified peptide ratios were quantified. Shown in red are probe-modified peptides that showed a ratio greater than 2 with adjusted p-values less than 0.01 with C619 of CTNNB1 shown as significantly engaged. **(c)** Cellular thermal shift assay with EN83 treatment. HiBiT-CTNNB1 HEK293 cells were treated with DMSO or EN83 (50 μM) for 1 h after which cells were heated to the designated temperatures, insoluble proteins were precipitated, and CTNNB1 and GAPDH were detected by Western blotting. **(d)** Quantification of experiment in **(c). (e)** Cysteine mutant rescue of EN83-mediated CTNNB1 degradation. HEK293T cells were transfected with FLAG-tagged wild-type (WT) or C439S, C466S, C520S, or C619S FLAG-tagged CTNNB1 mutants and treated with DMSO vehicle or EN83 (10 μM) for 24 h after which FLAG and loading control GAPDH were detected by Western blotting. **(f)** Quantification of experiment from **(e)**. Blots in **(a, c, e)** are representative of and quantitation of data in n=3 biologically independent replicates/group. Chemoproteomic data in **(b)** and quantified data in **(d, f)** are from n=3 biologically independent replicates/group. Data shown in **(d, f)** is shown as average ± sem values with individual replicates shown in **(f)**. Significance in **(d, f)** shown as *p<0.05 compared to DMSO treated controls for each temperature in **(d)** or each wild-type or mutant group in **(g)** and #p<0.05 compared to EN83-treated wild-type group in **(f)**.

We next sought to confirm the contributions of the cysteines targeted by EN83 in CTNNB1 leading to its degradation. We expressed either FLAG-tagged wild-type, C439S, C466S, C520S, or C619S mutant CTNNB1 in cells and showed that mutation of C466, C520, or C619, but not C439, to serines significantly attenuated CTNNB1 degradation in cells. These results demonstrated that direct targeting of three out of the four cysteines identified within CTNNB1 led to the degradation of CTNNB1.

### Improving Potency of EN83 Against CTNNB1

Towards improving potency of EN83 against CTNNB1 and exploring structure-activity relationships, we synthesized several analogs of EN83. EN83 is a racemic mixture of two enantiomers. We first separated the individual enantiomers, by chiral chromatographgy, to determine whether there was a particular enantiomer that was more potent. Both enantiomers of EN83 showed comparable lowering of CTNNB1 levels and showed equivalent CTNNB1 binding, indicating that there was no difference between the two enantiomers **(Figure S2a-S2c)**. We found that replacement of the methoxy groups with trifluoromethyl moieties, NF740 and NF741, was tolerated for both binding to CTNNB1 by gel-based ABPP and lowering CTNNB1 HiBiT levels in HEK293 cells in a dose-responsive manner **(Figure 4a-4c)**. This lowering of CTNNB1 was also confirmed to be proteasome-dependent by Western blotting **(Figure 4d)**. We also sought to replace the chloroacetamide cysteine-reactive warhead which may possess metabolic liabilities for further development. Replacement of the warhead with an acrylamide warhead found in many orally bioavailable covalent drugs, such as the BTK inhibitor ibrutinib or the KRAS G12C inhibitor sotorasib, unfortunately compromised binding to CTNNB1 and did not show lowering of CTNNB1 in either HiBiT or Western blotting detection approaches **(Figure S3a-S3d)**. We found that replacement of the chloroacetamide warhead with an oxo-phenylbutenamide warhead with NF764 and NF765 improved CTNNB1 binding and showed more potent loss of CTNNB1 HiBiT levels in cells **(Figure 4e-4g)**. We also confirmed that both NF764 and NF765 lowered CTNNB1 protein levels in cells in a proteasome-dependent manner **(Figure 4h)**. We thus demonstrated that the potency of EN83 can be improved through our preliminary medicinal chemistry efforts.

**Figure 4.**
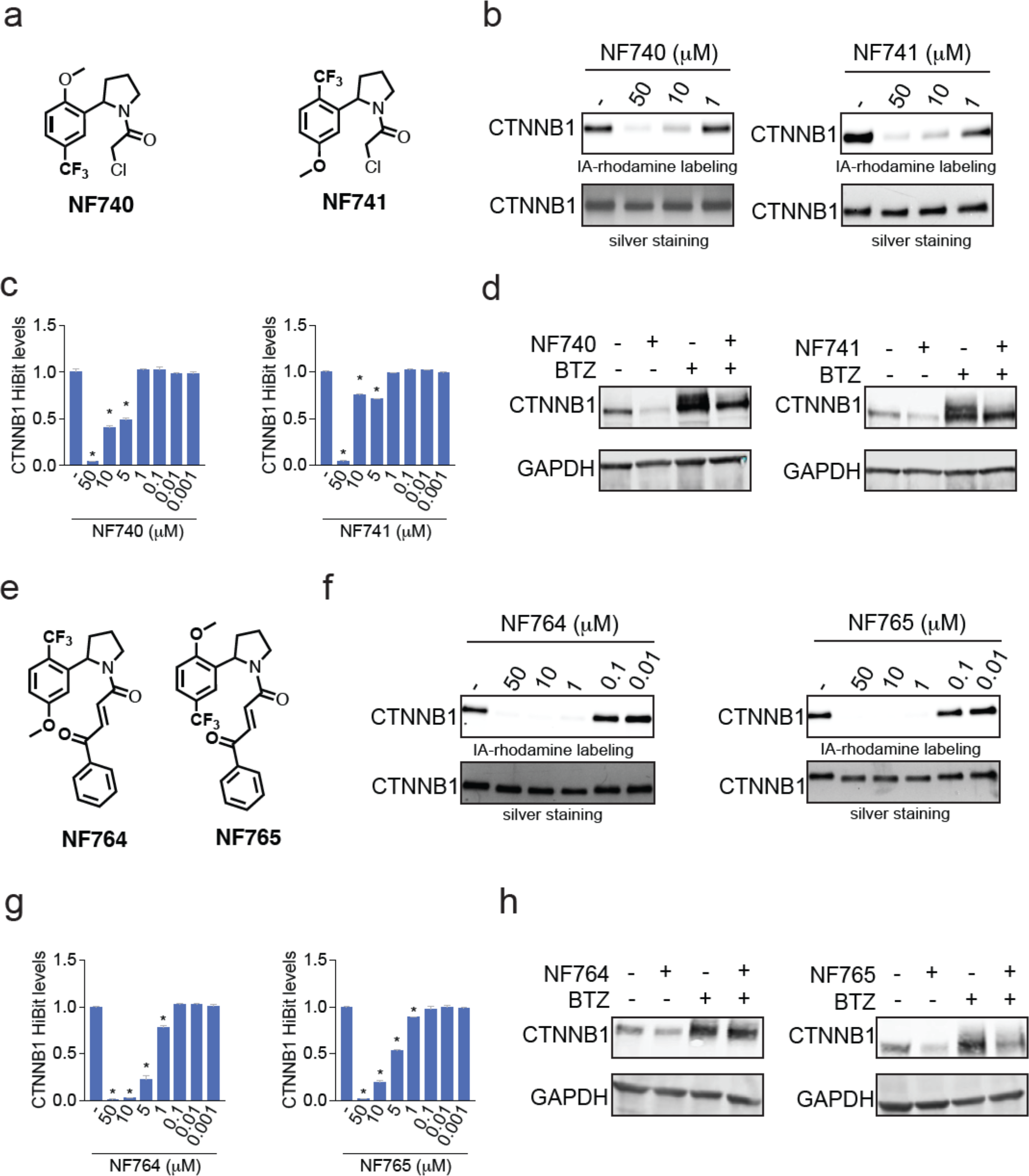
Structure-activity relationship of EN83 analogs. **(a)** Structures of EN83 analogs NF740 and NF741. **(b)** Gel-based ABPP analysis of EN83 against CTNNB1. CTNNB1 pure protein was pre-incubated with DMSO or compounds 30 min prior to IA-rhodamine labeling (100 nM) for 60 min, after which proteins were separated by SDS/PAGE and probe labeling was assessed by in-gel fluorescence and protein loading was assessed by silver staining. **(c)** HiBiT-CTNNB1 levels from NF740 or NF741 treatment. HiBiT-CTNNB1 HEK293 cells were treated with DMSO vehicle or compounds for 24 h. **(d)** Proteasome dependence of CTNNB1 loss. HiBiT-CTNNB1 HEK293 cells were pre-treated with DMSO vehicle or bortezomib (1 μM) for 1 h prior to treatment with DMSO vehicle or compound for 24 h. CTNNB1 and loading control GAPDH levels were assessed by Western blotting. **(e)** Structures of two additional more advanced analogs NF764 and NF765. **(f)** Gel-based ABPP of NF764 and NF765 using conditions described in **(b). (g)** HiBiT-CTNNB1 levels from NF764 and NF765 treatment using conditions described in **(c). (h)** Proteasome dependence of CTNNB1 loss using conditions described in **(d)**. Gels and blots in **(b, d, f, h)** are representative of n=3 biologically independent replicates/group. Bar graphs in **(c, g)** shown as average ± sem values. Significance in **(c, g)** shown as *p<0.05 compared to vehicle treated controls.

## Conclusions

In this study, we put forth a covalent ligand degrader of CTNNB1 that acts through direct covalent targeting of three distinct cysteines in the armadillo repeat domains of CTNNB1—C466, C520, and C619—to destabilize and degrade CTNNB1 in a proteasome-dependent manner. We conjecture that the mode of degradation observed in this study is likely distinct from traditional modes of targeted protein degradation, such as Proteolysis Targeting Chimeras (PROTACs) and molecular glue degraders; and instead occurs through a thermodynamic destabilization of CTNNB1 resulting from direct covalent binding, leading to a destabilization-mediated degradation of CTNNB1 ^19–21^. We believe this is analogous to our previous findings with another oncogenic transcription factor MYC, where we discovered a covalent ligand EN4 that irreversibly targeted C171 in MYC leading to a thermal destabilization and loss of MYC protein levels ^15^. Many transcription factors have often eluded classical drug discovery efforts because of their lack of well-defined binding pockets for small-molecule binding and also because many of them possess large segments of unstructured regions. Covalent targeting of ligandable cysteines within transcription factors causing their destabilization and degradation may represent a potential strategy for pharmacologically tackling this challenging class of targets.

While we demonstrate initial medicinal chemistry efforts to explore structure-activity relationships of EN83 leading to improved CTNNB1 binders and degraders, further medicinal chemistry efforts need to be performed to further improve the potency, selectivity, and metabolic stability of the compounds reported here. We note that while we observed CTNNB1 loss at 24 h with EN83 treatment in the model cell line HEK293 cells, we only observed acute loss of CTNNB1 at 2 and 4 h with recovery by 24 h in more relevant colorectal cancer cell lines. CTNNB1 is heavily regulated by the ubiquitin-proteasome system and thus may have faster turnover in cancer cell lines that are driven by CTNNB1^2^. As such, a covalent degrader of CTNNB1 would likely need to maintain target engagement and ultimately show sustained *in vivo* pharmacokinetics to achieve persistent pharmacodynamics. A covalent degrader that irreversibly engages CTNNB1 will show stoichiometric and non-catalytic degradation. A covalent reversible CTNNB1 degrader may be able to exploit the ligandable cysteines within CTNNB1 while achieving sub-stoichiometric catalytic degradation of CTNNB1.

Overall, our study demonstrates the utility of using covalent chemoproteomic strategies to identify unique ligandable sites and covalent ligands that can directly target classically intractable target classes such as transcription factors to ultimately degrade targets such as CTNNB1 for potential future cancer therapy applications.

## Acknowledgement

We thank the members of the Nomura Research Group and Bristol Myers Squibb for critical reading of the manuscript. This work was supported by Bristol Myers Squibb for all listed authors. This work was also supported by the Nomura Research Group and the Mark Foundation for Cancer Research ASPIRE Award for DKN. This work was also supported by grants from the National Institutes of Health (R01CA240981 and R35CA263814 for DKN). We also thank Dr. Hasan Celik and UC Berkeley’s NMR facility in the College of Chemistry (CoC-NMR) for spectroscopic assistance. Instruments in the College of Chemistry NMR facility are supported in part by NIH S10OD024998.

## Author Contributions

AG, NF, IEW, DKN conceived of the project idea. AG, NF, DKN designed experiments, performed experiments, analyzed and interpreted the data, and wrote the paper. OWH performed experiments and provided protein reagents. AG, NF, JH, YW, BEAP, DKN performed experiments, analyzed and interpreted data, and provided intellectual contributions.

## Methods

### Cell culture

CTNNB1 HiBiT cells were commercially purchased through Promega (CS302340). HEK293, COLO-201, HT29, and SW480 were obtained from UC Berkeley’s Biosciences Divisional Services Cell Culture Facility. HiBiT, HEK, and HT29 were cultured in DMEM and COLO-201 cells were cultured in RPMI base media. Both media types contained 10% (v/v) fetal bovine serum (FBS), were supplemented with 1% glutamine, and were maintained at 37°C with 5% CO_2_.

### Covalent Ligand Library and Synthesis of Other Compounds

Covalent ligands starting with “EN” are commercially available from Enamine LLC. Synthesis of other compounds are described in **Supporting Information**.

### Covalent Ligand screen with CTNNB1 HiBiT Cell Line

Covalent ligand screen and dose responses were conducted using Promega Nano-Glo HiBiT Lytic Detection System (N3040) and CellTiter-Glo 2.0 Assay (G9242). HiBiT cells were seeded into 96-well plates (Corning 3917) at 35,000 cells per 100μL of media and were left overnight to adhere. Cells were treated with 25μL of media containing 1:125 dilution from a 1000x DMSO compound stock and treated for either 6hrs, 12hrs, or 24hrs. The lytic detection system recipe was followed per Promega’s suggestion. 125 μL of CTG or Lytic detection system reagents were added to each well. To assess the cells in the supernatant, half of the plates had the media removed prior to the addition of CTG or lytic detection system reagents. Plates were rocked for 15 minutes prior to their luminescence readout on the Tecan Spark Plate reader (30086376).

### Western Blotting

Pelleted cells are lysed with CST buffer containing protease inhibitor cocktail (Pierce A32955). Samples were centrifuged at 20,000 g for 20 min at 4 °C to remove cell debris. Samples’ protein content was normalized to run 30μg per well. Samples were then boiled for 8 min at 90 °C after addition of 4×reducing Laemmli SDS sample loading buffer (Alfa Aesar) and ran on precast 4−20% Criterion TGX gels (Bio-Rad).

Antibodies to CTNNB1 (Cell Signaling Technology, 8814S), GAPDH (Proteintech Group Inc., 60004–1-Ig or Cell Signaling Technology, 14C10), and DDDDK tag (Abcam, ab205606) were diluted per recommended manufacturers’ procedures. Proteins were resolved by SDS/PAGE and transferred to nitrocellulose membranes using the BioRad system (1704271 and 1704150). Blots were blocked with 5 % BSA in Tris-buffered saline containing Tween 20 (TBST) solution for 1 hour at room temperature, washed in TBST, and probed with primary antibody diluted in recommended diluent per manufacturer overnight at 4°C. Following washes with TBST, the blots were incubated in the dark with secondary antibodies purchased from Ly-Cor and used at 1:10,000 dilution in 5% BSA in TBST at room temperature. Blots were visualized using an Odyssey Li-Cor scanner after additional washes. If additional primary antibody incubations were required the membrane was stripped using ReBlot Plus Strong Antibody Stripping Solution (EMD Millipore, 2504), washed and blocked again before being reincubated with primary antibody.

### β‐Catenin Constructs

Human Beta catenin (residues 151-661) constructs were synthesized and inserted into a pET28a vector. For synthesis of the construct, a six-histidine tag next to a TEV protease site were placed on the n-terminal side of Beta catenin and a two-repeat linker of AEAAA followed by human TF7L2 18-49/avi on the c-terminal end of Beta catenin.

### β-Catenin Purification

Beta catenin constructs were expressed in *E*.*coli* DE3 pLys cells grown to OD of 600 at 37 °C and induced with 300 μM IPTG. Expression was allowed to proceed for 16 hr at 16 °C. Cells were centrifuged at 5000 x g for 10 min then collected and frozen at -80 °C. *E. coli* cells were lysed three passes through a fluid homogenizer packed in ice. Cell debris was centrifuged at 103,000 x g for 30 min, supernatant was added to 4 ml of Talon and allowed to batch bind for 6 0min at 4 °C. Talon resin was washed with 50 CV of 50 mM TRIS pH 8.5, 500 mM NaCl, 10% glycerol, 0.5 mM TCEP and 25 mM imidazole. Protein was eluted with 5 0mM TRIS pH 8.5, 500 mM NaCl, 10% glycerol, 0.5 mM TCEP and 300 mM imidazole until no protein could be visually detected by Coomassie blue protein assay. Eluted Beta catenin avi tagged protein was biotinylated with BirA enzyme overnight. Biotinylated protein was desalted to remove any free biotin then allowed to batch bind for 2 hrs to overnight to Streptavidin Mutein matrix. The protein was then eluted with 50 mM TRIS pH 8.5, 500 mM NaCl, 10% glycerol, 0.5 mM TCEP and 2 mM biotin. Eluted protein was directly run over a S200 16/600 SEC column (50 mM TRIS pH 8.5, 500 mM NaCl, 10% glycerol, 0.5 mM TCEP).

### Covalent Ligand screen and Gel-Based ABPP with CTNNB1 Pure Protein

CTNNB1 pure protein (0.1μg/25 μL in PBS) was treated with either DMSO vehicle or covalent ligand at 37 °C for 30 min, and subsequently treated with 0.1 μM IA-Rhodamine (Setareh Biotech) for 1 h at RT in the dark. The reaction was stopped by addition of 4×reducing Laemmli SDS sample loading buffer (Alfa Aesar). After boiling at 95 °C for 5 min, the samples were separated on precast 4−20% Criterion TGX gels (Bio-Rad) and were analyzed by in-gel fluorescence using a ChemiDoc MP (Bio-Rad).

### NF686 Probe labeling on CTNNB1 NV03 Pure Protein

CTNNB1 pure protein (0.2μg/50 μL in PBS) was treated with either DMSO vehicle or NF686 at 37°C for 30min. Click reagents Azide-Fluor 545 (Click Chemistry Tools, Inc. AZ109-5), Copper (II) Sulfate, and TBTA (TCI Chemicals, T2993) were added to have a final concentration of 21.8 μM, 873.4 μM, and 47.2 μg/mL, respectively, for 1hr at RT in the dark. The reaction was stopped by addition of 4×reducing Laemmli SDS sample loading buffer (Alfa Aesar). After boiling samples at 95 °C for 5 min, the samples were separated on precast 4−20% Criterion TGX gels (Bio-Rad). Probe-labeled proteins were analyzed by in-gel fluorescence using a ChemiDoc MP (Bio-Rad).

### Luciferase Reporter Assay

HEK293 cells were seeded at a density of 30,000/well in 96-well white plate overnight. Each well was transiently co-transfected with 60ng TCF/LEF luciferase reporter vector and 60 ng renilla luciferase control vector using Lipofectamine 2000, provided by TCF/LEF Reporter Kit Wnt / β-catenin signaling pathway (BPS Bioscience, #60500). After 72hrs post-transfection, cells were then treated with EN83 or DMSO vehicle control until desired time point. Firefly and renilla luciferase reporter genes’ expression levels were examined by TWO-Step Luciferase (Firefly & Renilla) Assay System (BPS Bioscience #60683) according to manufacturer’s protocols. Luminescent signals were measured using the Tecan Spark Plate reader (30086376).

### Chemoproteomic Profiling

Chemoproteomic profiling methods are described in **Supporting Information**.

