## Supporting Information for "Covalent Degrader of the Oncogenic Transcription Factor β-Catenin"

**Supporting Table Legends**

**Table S1. Structures of compounds screened and screening data.** Tab 1 contains the structures of the cysteine-reactive covalent ligands screened in this study. Tab 2 contains the covalent ligand screening data. HEK293 cells expressing a N-terminal HiBiT tagged CTNNB1 in the endogenous CTNNB1 locus were treated with DMSO vehicle or a cysteine-reactive covalent ligand (50 μM) for 24 h and HiBiT-CTNNB1 levels were quantified. Reported are the compound/control ratio values.

**Table S2. Cysteine chemoproteomic profiling of EN83 by isoDTB-ABPP**. HiBiT-CTNNB1 HEK293 cells were treated with DMSO vehicle or EN83 (10 μM) for 2 h. Lysates were then labeled with an alkyne-functionalized iodoacetamide probe (IA-alkyne) (200 μM) for 1 h, followed by appendage of isotopically light or heavy desthiobiotin-azide handles by CuAAC, after which probe-modified proteins were avidin-enriched, tryptically digested, and probe-modified peptides were eluted and analyzed by LC-MS/MS and control versus treated or light versus heavy probe-modified peptide ratios were quantified. Shown in the first tab is the totality of all probe-modified peptides detected. Shown in the second tab are the probe-modified peptides detected in all 3 biologically independent replicates and their quantification, statistical significance, peptide sequence, site of probe modification, individual ratio replicates, and protein identification.

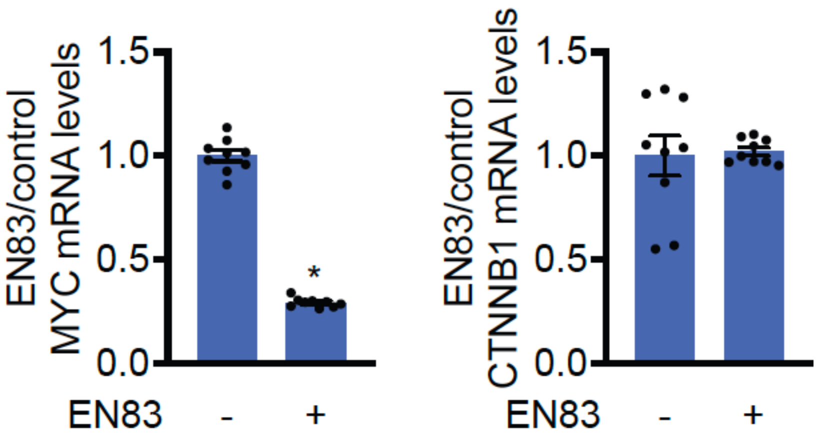


**Figure S1. CTNNB1 and CTNNB1 gene target MYC mRNA levels.** HEK293 cells were treated with DMSO vehicle or EN83 (10 μM) for 24 h and CTNNB1 and MYC mRNA levels were assessed by qPCR. Data shown are average ± sem values from n=9 biologically independent replicates/group. Significance is expressed as *p<0.05 compared to vehicle-treated controls.

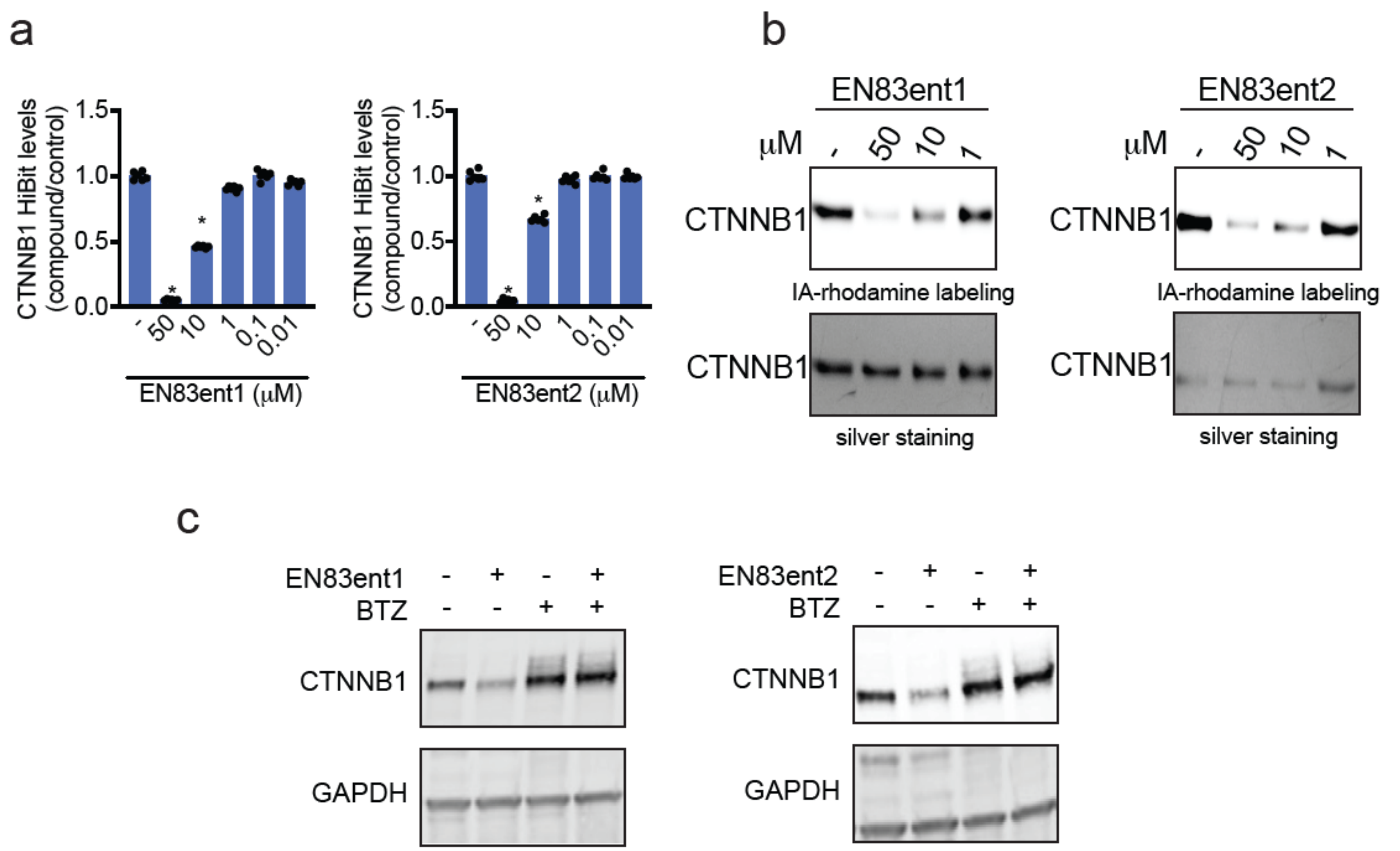


**Figure S2. Characterization of EN83 enantiomers. (a)** HiBiT-CTNNB1 levels from treatment of cells with EN83 enantiomers. HiBiT-CTNNB1 HEK293 cells were treated with DMSO vehicle or compounds for 24 h. **(b)** Gel-based ABPP analysis of EN83 enantiomers against CTNNB1. CTNNB1 pure protein was pre-incubated with DMSO or compounds 30 min prior to IA-rhodamine labeling (100 nM) for 60 min, after which proteins were separated by SDS/PAGE and probe labeling was assessed by in-gel fluorescence and protein loading was assessed by silver staining. **(c)** Proteasome dependence of CTNNB1 loss. HiBiT-CTNNB1 HEK293 cells were pre-treated with DMSO vehicle or bortezomib (1 μM) for 1 h prior to treatment with DMSO vehicle or compound for 24 h. CTNNB1 and loading control GAPDH levels were assessed by Western blotting. Gels and blots in **(b, c)** are representative of n=3 biologically independent replicates/group. Bar graphs in **(a)** shown as average ± sem values. Significance in **(a)** shown as *$p<0.05$ compared to vehicle treated controls.

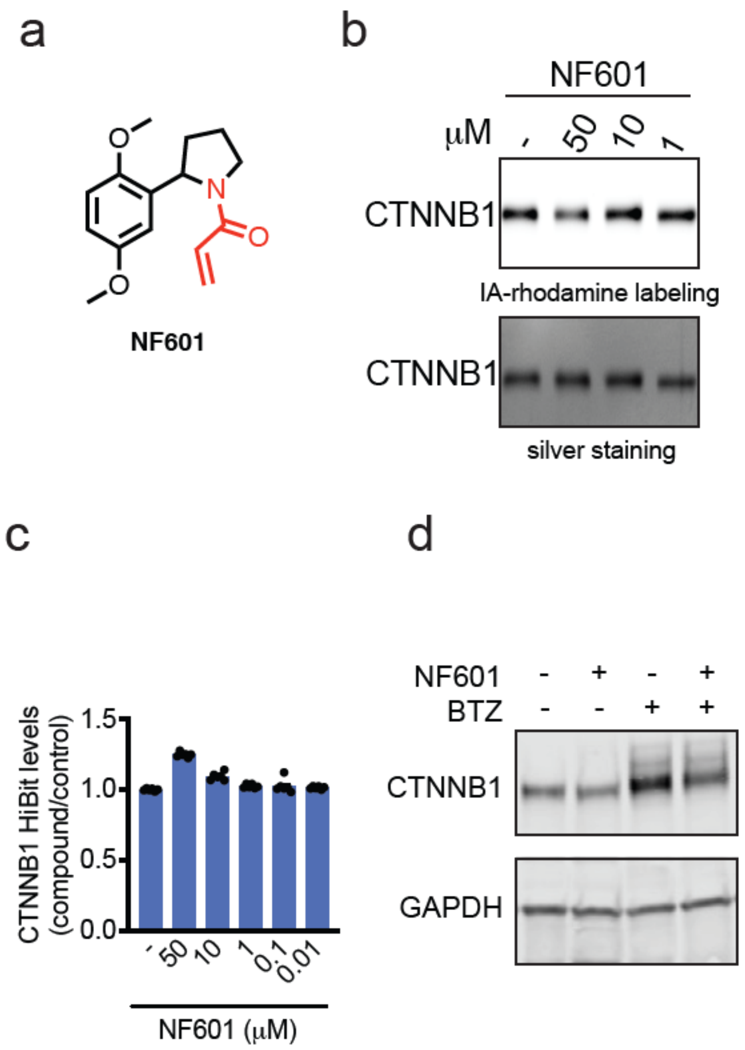


**Figure S3. Characterization of NF601. (a)** Structure of EN83 analog with acrylamide warhead, NF601. **(b)** Gel-based ABPP analysis of NF601 against CTNNB1. CTNNB1 pure protein was pre-incubated with DMSO or NF601 30 min prior to IA-rhodamine labeling (100 nM) for 60 min, after which proteins were separated by SDS/PAGE and probe labeling was assessed by in-gel fluorescence and protein loading was assessed by silver staining. **(c)** HiBiT-CTNNB1 levels from treatment of cells with EN83 enantiomers. HiBiT-CTNNB1 HEK293 cells were treated with DMSO vehicle or NF601 for 24 h. **(d)** Proteasome dependence of CTNNB1 loss. HiBiT-CTNNB1 HEK293 cells were pre-treated with DMSO vehicle or bortezomib (1 μM) for 1 h prior to treatment with DMSO vehicle or NF601 for 24 h. CTNNB1 and loading control GAPDH levels were assessed by Western blotting. Gels and blots in **(b, d)** are representative of n=3 biologically independent replicates/group. Bar graphs in **(c)** shown as average ± sem values.

**Supporting Information**

**Bortezomib or MLN4924 Rescue Studies**

For 96-well plate format, HiBiT cells were plated at 35,000 cells per 100μL of media and were left overnight to adhere. Cells were pretreated for 1hr with 12.5μL of media containing Bortezomib (Cayman, C835F70) or MLN4924 (Tocris Bioscience, 649910) at a final concentration of 1μM. 12.5μL of media containing covalent ligand (1:62.5) was then added and incubated for 24hrs. 125 μL of Lytic detection system reagent were added to each well. To assess the cells in the supernatant, half of the plates had the media removed prior to the addition of CTG or lytic detection system reagents. Plates were rocked for 15 minutes prior to their luminescence read on the Tecan Spark Plate reader (30086376).

For Western blots, 8E6 cells per 10mL of media were plated in 10cm plates and left overnight to adhere. Cells were pretreated for 1hr with either Bortezomib (Cayman, C835F70) or MLN4924 (Tocris Bioscience, 649910) at a final concentration of 1μM. Cells were then treated with EN83 until desired time point. Cells from both the supernatant and on the plate were harvested and assessed via western blot.

**Cellular Thermal Shift Assay (CETSA)**

HiBiT CTNNB1 cells were seeded at a density of 8E6 cells per 10mL of media and were treated with either DMSO vehicle control or EN83 for 1hr. Cells were harvested and washed twice with PBS, then suspended in 1 mL of PBS containing protease inhibitor cocktail (Pierce A32955). The cell suspension was allocated into eight 0.2 mL PCR tubes with 100μL volume per tube. PCR strips were designated a temperature via a gradient program on Bio-Rad's T100 Thermal cycler. Samples were heated at their respective temperatures (52.0 °C, 53.9 °C, 56.8 °C, 61.3 °C, 66.8 °C, 71.9 °C, 75.1 °C, and 77.0 °C) for 3 min, then at 25 °C for 3 min. Afterwards, cells were immediately snap-lysed in liquid nitrogen (3 freeze-thaw cycles). Cell debris along with precipitated and aggregated proteins were removed by centrifuging samples at 20,000 g for 20 min at 4 °C. 80μL of centrifuged samples were transferred to new PCR tubes and boiled for 8 min at 90 °C after addition of 4×reducing Laemmli SDS sample loading buffer (Alfa Aesar). Samples were analyzed by Western Blot analysis. Protein intensity was quantified through Image J software.

**CST Lysis Buffer Recipe**

Working lysis buffer is made with 1 tablet of protease inhibitor cocktail (Pierce A32955) solubilized in 1mL PBS and 9mL of CST buffer.

| | |
|---|---|
| Cold MilliQ Water | 70mL |
| 1M Tris, pH 7.5 | 2mL |
| 5M NaCl | 3mL |
| 5% Triton X-100 | 20mL |
| 1M NaF | 5mL |
| 0.5M EDTA, pH 8.0 | 200μL |
| 0.5M EGTA, pH 8.0 | 200μL |
| 200mM Sodium pyrophosphate | 1250μL |
| 1M -glycerophosphate | 200 μL |
| 100mM OVO4 | 1mL |

**Gene Expression RTqPCR**

Total RNA was extracted from cells using Trizol (Thermo Fisher Scientific) and aqueous solution was processed using QIAgen RNeasy Mini Kit (catalogue no. 74104). cDNA was synthesized using Qiagen Quantitect Reverse

Transcription (205311) and gene expression was confirmed by qPCR using the manufacturer's protocol Power SYBR Green Master Mix 2X concentration (4368577) on the CFX Connect Real-Time PCR Detection System (BioRad). Primer sequences for SYBR Green were derived from Primer Bank. Sequences of primers are as follow:

GAPDH Rev: GGACCTGACCTGCCGTCTAG
GAPDH Rev: TAGCCCAGGATGCCCTTGAG
CTNNB1 Fwd: CAC AAG CAG AGT GCT GAA GGT G
CTNNB1 Rev: GAT TCC TGA GAG TCC AAA GAC AG
cMYC Fwd: TGCTGCCAAGAGGGTCAAG
cMYC Rev: GCGCTCCAAGACGTTGTGTGT

**Site-Directed Mutagenesis on FLAG-tagged wild type CTNNB1 plasmid**
Original CTNNB1 plasmid was purchased from Origene (RC208947) and site-directed mutagenesis were performed using Agilent's QuikChange Lightning Site-Directed Mutagenesis Kit (210518). Thermal cycle time & temperature, DpnI digestion, and transformation conditions were followed as suggested by user guide. Mutant plasmids were initially transformed into XL10-Gold Ultracompetent cells provided by Agilent. Plasmids were then isolated and re-transformed into a high-plasmid yielding E. coli background: Stbl3 (C7373-03). Transformation conditions suggested by Agilent or Invitrogen were followed.

The following primer pairs were designed using Agilent's web-based primer design tool:

**C619S Primer #1:** CCT GAG CAA GTT CAC TGA GGA CCC CTG CAG
**C619S Primer #2**: CTG CAG GGG TCC TCA GTG AAC TTG CTC AGG

**C466S Primer #1:** GAT GAC GAA GAG CAC TGA TGG CAG GCT CAG T
**C466S Primer #2**: ACT GAG CCT GCC ATC AGT GCT CTT CGT CAT C

**C520S Primer #1:** TGA TTT GCG GGA CTA AGG GCA AGA TTT CGA ATC AAT CC
**C520S Primer #2**: GGA TTG ATT CGA AAT CTT GCC CTT AGT CCC GCA AAT CA

**C439S Primer #1:** TAT ACC ACC CAC TTG GCT GAC CAT CAT CTT GTT CT
**C439S Primer #2**: AGA ACA AGA TGA TGG TCA GCC AAG TGG GTG GTA TA

**Media and Agar Plate Preparation for Site-Directed Mutagenesis**
NZY+ Broth recipe suggested by Agilent's Site-Directed Mutagenesis kit were followed. LB Broth Miller (BP1426-500) recipe suggested by Fisher Bioreagents were followed.

LB agar plates were made with BD Difco Dehydrated Culture Media: LB Agar, Lennox (240110). Measurements and autoclaving conditions suggested by manufacturer were followed. After agar cooled to 50°C, LB agar was spiked with 25 µg/ml Kanamycin, 80 µg/ml X-gal and 20 mM IPTG and poured into plates.

**Bacterial Culture**
Plasmids transformed in XL10-Gold Ultracompetent cells were cultured in NZY+ broth with Kanamycin overnight at 200rpm at 37C**.** The commercially available plasmid and in-house produced mutants that were transformed into Stbl3 competent cells were cultured in LB with Kanamycin overnight at 200rpm at 37C**.**

**Plasmid Isolation**

E. coli containing desired plasmids were pelleted, lysed, and neutralized using commercially available QIAprep Spin Miniprep's (catalogue no. 27104) user manual. The plasmid elutes' concentration were determined using Nanodrop quantification.

**Primers for Mutant Plasmid Sequence Confirmation**

Plasmids were sent to Elim Biopharm to confirm mutant sequences. The following primers were used:

VP1.5: GGACTTTCCAAAATGTCG
XL39: ATTAGGACAAGGCTGGTGGG
CTNNB1 Mutant Primer #1: GAT GCA GAA CTT GCC ACA CG
CTNNB1 Mutant Primer #2: GGA GCT AAA ATG GCA GTG CG

**Expression of FLAG-tagged wild type CTNNB1 and FLAG-tagged C439S, C466S, C520S, C619S mutants**

FLAG-tagged CTNNB1 plasmids (WT Origene Technologies Inc., RC208947), C439S, C466S, C520S, and C619S plasmids were transfected into HEK293T cells. HEK293T cells were grown to 30-50% confluency. Immediately before transfection, media was replaced with fresh DMEM. Each plate was transfected with 24 µg of overexpression plasmid and with 24 µL Lipofectamine 2000 (ThermoFisher, 11668027) in Opti-MEM (ThermoFisher, 31985062). After 48 h, the media was changed to fresh DMEM and cells were treated with either DMSO or EN83 (10 µM) for 24 h. Cells were then harvested, and protein abundance was analyzed by Western blotting, using Anti-DDDDK tag (Abcam, ab205606) or GAPDH (Cell Signaling Technology, 14C10) antibodies.

**Pulldown of CTNNB1 from HEK293T-CTNNB1 HiBiT Cells with NF686 probe**

HEK293T-CTNNB1 HiBiT cells were treated at 70% confluency with bortezomib (1 µM) for 1h, followed by DMSO or NF686 (50 µM) for 24 h. Cells were harvested, lysed via sonication (a hard-spin was not performed) and the resulting lysate normalized to 5 mg/mL per sample. 500 µL of each lysate sample was incubated for 1 h at RT with 10 µL of 10 mM biotin picolyl azide (in DMSO) (Sigma Aldrich 900912), 10 µL of 50 mM TCEP (in $H_2O$), 30 µL of TBTA ligand (0.9 mg/mL in 1:4 DMSO/tBuOH), and 10 µL of 50 mM $CuSO_4$. Proteins were precipitated, washed 3 × with cold MeOH, resolubilized in 200 µL of 1.2% SDS/PBS (w/v) and heated for 5 min at 90 °C. 10 µL of each sample was removed for Western Blot analysis of input. To the remaining 190 µL was added 1 mL PBS and 70 µL streptavidin agarose beads (ThermoFisher, 20353). Samples were incubated at 4 °C overnight on a rotator. The following day the samples were warmed to RT and washed with 0.2% SDS and further washed 3 × with 500 µL PBS and 3 × with 500 µL $H_2O$ to remove non-probe-labeled proteins. The washed beads were resuspended in 30 µL Laemmli SDS sample loading buffer (Alfa Aesar), heated to 95 °C for 5 min and analyzed by Western Blot.

**IsoDTB-ABPP Cysteine Chemoproteomic Profiling of EN83**

HEK293T-CTNNB1 HiBiT cells were treated with either EN83 (10 µM) or DMSO for 2 h before cell collection and lysis. The proteome concentrations were determined using BCA assay and adjusted to 2 mg/mL. For each biological replicate, 2 aliquots of 1 mL of 2 mg/mL were used (i.e. 4 mg per condition). Each aliquot was treated with 20 µL of IA-alkyne (26.6 mg/mL in DMSO, 200 µM final concentration) for 1 h at RT. Two master mixes of the click reagents were prepared in the meanwhile, each containing 1020 µL TBTA (0.9 mg/mL in 4:1 tBuOH/DMSO), 330 µL $CuSO_4$ (12.5 mg/mL in $H_2O$), 330 µL TCEP (14.0 mg/mL in $H_2O$) and 160 µL of either heavy or light isoDTB tags (4 mg in DMSO, Click Chemistry Tools, 1565). The samples were then treated with 120 µL of the heavy (DMSO treated) or light (compound treated) master mix for 1 h at RT. After incubation, one light and one heavy-labeled samples were combined and acetone-precipitated overnight at -20 °C. The samples were then centrifuged at 3,500 rpm for 10 min, acetone was removed and they were resuspended in cold MeOH by sonication. They were centrifuged at 3,500 rpm for 10 min and MeOH was removed (repeated 3× in total). The pellets were dissolved in 300 µL urea (8M in 0.1 M TEAB) by sonication and the urea concentration was then adjusted to 2M by adding 900 µL of TEAB (0.1 M). Two tubes containing solubilized proteins were combined, further diluted with 2400 μL 0.2% NP40 in PBS and bound to high-capacity streptavidin agarose beads (250 μL/sample, ThermoFisher, 20357) for 1 h at RT with mixing. The beads were then centrifuged for 1 min at 1,000 g, the supernatant was removed and the beads were washed 3 times with 0.1% NP40 in PBS, 3 times with PBS and 3 times with $H_2O$. They were then resuspended in 8M urea (600 μL in 0.1 M TEAB) and treated with DTT (30 μL, 31 mg/mL in $H_2O$) for 45 min at 37 °C. They were then reacted with iodoacetamide (30 μL, 74 mg/mL in $H_2O$) for 30 min at RT, followed by DTT (30 μL, 31 mg/mL in $H_2O$) for 30 min at RT. The samples were diluted with 1800 μL TEAB (0.1 M), centrifuged for 1 min at 1,000 g and the supernatant was removed. The beads were resuspended in 400 μL urea (2M in 0.1 M TEAB), trypsin (8 μL, 0.5 mg/mL) was added, followed by 1 μL $CaCl_2$ (100 mM in $H_2O$) and the samples incubated for 19 h at 37 °C. The samples were then diluted with 800 μL 0.1% NP40 in PBS and the beads were washed 3 times with 0.1% NP40 in PBS, 3 times with PBS and 3 times with $H_2O$. Peptides were then eluted with 0.1% formic acid in 50% acetonitrile (3 × 400 μL). The samples were then dried by using a vacuum concentrator at 30 °C, resuspended in 300 μL 0.1% TFA in $H_2O$, and fractionated using high pH reversed-phase peptide fractionation kits (ThermoFisher, 84868) according to manufacturer's protocol.

**IsoDTB Mass Spectrometry Analysis**

Mass spectrometry analysis was performed on an Orbitrap Eclipse Tribrid Mass Spectrometer with a High Field Asymmetric Waveform Ion Mobility (FAIMS Pro) Interface (Thermo Scientific) with an UltiMate 3000 Nano Flow Rapid Separation LCnano System (Thermo Scientific). Off-line fractionated samples (5 μl aliquot of 15 μl sample) were injected via an autosampler (Thermo Scientific) onto a 5 μl sample loop which was subsequently eluted onto an Acclaim PepMap 100 C18 HPLC column (75 μm x 50 cm, nanoViper). Peptides were separated at a flow rate of 0.3 μl/min using the following gradient: 2 % buffer B (100 % acetonitrile with 0.1 % formic acid) in buffer A (95:5 water:acetonitrile, 0.1 % formic acid) for 5 min, followed by a gradient from 2 to 40 % buffer B from 5 to 159 min, 40 to 95 % buffer B from 159 to 160 minutes, holding at 95 % B from 160-179 min, 95 % to 2 % buffer B from 179 to 180 min, and then 2 % buffer B from 180 to 200 min. Voltage applied to the nano-LC electrospray ionization source was 2.1 kV. Data was acquired through an MS1 master scan (Orbitrap analysis, resolution 120,000, 400-1800 m/z, RF lens 30 %, heated capillary temperature 250 °C) with dynamic exclusion enabled (repeat count 1, duration 60 s). Data-dependent data acquisition comprised a full MS1 scan followed by sequential MS2 scans based on 2 s cycle times. FAIMS compensation voltages (CV) of -35, -45, and -55 were applied. MS2 analysis consisted of: quadrupole isolation window of 0.7 m/z of precursor ion followed by higher energy collision dissociation (HCD) energy of 38 % with a orbitrap resolution of 50,000.

Data were extracted in the form of MS1 and MS2 files using Raw Converter (Scripps Research Institute) and searched against the Uniprot human database using ProLuCID search methodology in IP2 v.3-v.5 (Integrated Proteomics Applications, Inc.) [1]. Cysteine residues were searched with a static modification for carboxyaminomethylation (+57.02146) and up to two differential modifications for methionine oxidation and either the light or heavy isoDTB tags (+561.33872 or +567.34621, respectively). Peptides were required to be fully tryptic peptides. ProLUCID data were filtered through DTASelect to achieve a peptide false-positive rate below 5%. Only those probe-modified peptides that were evident across two out of three biological replicates were interpreted for their isotopic light to heavy ratios. Light versus heavy isotopic probe-modified peptide ratios are calculated by taking the mean of the ratios of each replicate paired light versus heavy precursor abundance for all peptide-spectral matches associated with a peptide. The paired abundances were also used to calculate a paired sample *t*-test *P* value in an effort to estimate constancy in paired abundances and significance in change between treatment and control. *P* values were corrected using the Benjamini–Hochberg method.

**Synthetic Methods and Characterization**

All chemical reactions were carried out under a nitrogen atmosphere with dry solvents under anhydrous conditions, unless otherwise noted. Reagents were purchased at the highest commercial quality and used without further purification, unless otherwise stated. Room temperature is defined as between 21-25 °C. Reactions were stirred magnetically and monitored by thin layer chromatography (TLC) using TLC plates pre-coated with silica gel 60 F254 on aluminium (Merck KGaA). Detection was by UV (254 nm and 365 nm) or chemical stain ($KMnO_4$, ninhydrin, iodine). Solvents were removed *in vacuo* using a Buchi R-300 Rotavapor (equipped with an I-300 Pro Interface, B-300 Base Heating Bath, Welch 2037B-01 DryFast pump, and VWR AD15R-40-V11B Circulating Bath). Solvents for silica gel chromatography were used as supplied by Sigma-Aldrich. Automated flash chromatography was performed on a Biotage Isolera instrument, equipped with a UV detector. Chromatograms were recorded at 254 and 280 nm. High-resolution mass spectra (HRMS) were obtained on a Q Exactive Plus mass spectrometer (Thermo Fisher Scientific). $^1$H and $^{13}$C Nuclear Magnetic Resonance (NMR) spectra were recorded on BRUKER AV spectrometer operating at 600 MHz for $^1$H and at 150 MHz for $^{13}$C NMR. Measurements were carried out at ambient temperature. Chemical shifts (δ) are reported in ppm with the residual solvent signal as internal standard (chloroform at 7.26 and 77.2 ppm for $^1$H NMR and $^{13}$C NMR, respectively). The multiplicity of each signal is indicated as s = singlet, d = doublet, t = triplet, q = quartet, quin = quintet, m = multiplet (i.e. complex peak obtained due to overlap), app = apparent or a combination of these. Coupling constants (J) are reported in Hertz (Hz). $^{13}$C NMR spectra were recorded with broadband $^1$H decoupling.

**1-(2-(2,5-dimethoxyphenyl)pyrrolidin-1-yl)prop-2-en-1-one (NF601)**

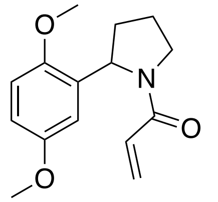

2-(2,5-Dimethoxyphenyl)pyrrolidine (100 mg, 0.483 mmol) was dissolved in DMF (10 mL). To this was added acryloyl chloride (94.0 µL, 1.16 mmol), followed by $NEt_3$ (241 µL, 1.73 mmol) and the resultant mixture was stirred at RT for 18 h. It was then concentrated *in vacuo* and purification by column chromatography (gradient elution from $CH_2Cl_2$ to 70% diethyl ether in $CH_2Cl_2$) afforded the target product as a clear oil (89.0 mg, 0.341 mmol, 71%).

$^1$H NMR ($CDCl_3$, 600 MHz) $\delta_H$ 6.82-6.67 (m, 2H), 6.59-6.52 (m, 1H), 6.37-6.29 (m, 1H), 6.12-6.07 (m, 1H), 5.99-5.28 (m, 2H), 3.85-3.81 (m, 4H), 3.74-3.68 (m, 4H), 2.35-2.19 (m, 1H), 1.97-1.80 (m, 3H); $^{13}$C NMR ($CDCl_3$, 150 MHz) $\delta_C$ 165.2, 164.1, 153.7, 153.4, 150.5, 149.8, 132.6 132.3, 128.9, 128.7, 127.9, 127.3, 113.2, 113.0, 112.0, 111.4, 111.1, 110.6, 56.5, 56.4, 55.9, 55.7, 55.7, 55.6, 47.7, 47.0, 34.1, 32.2, 32.7, 21.7; HRMS (ES+) calcd for $C_{15}H_{19}NO_3Na$ $[M+Na]^+$ 284.1263, observed 284.1252.

**1-(2-(2,5-dimethoxyphenyl)pyrrolidin-1-yl)propan-1-one (NF602)**

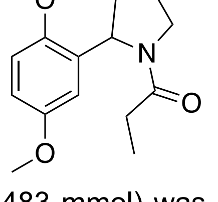

2-(2,5-dimethoxyphenyl)pyrrolidine (100 mg, 0.483 mmol) was dissolved in DMF (10 mL). To this was added propionyl chloride (101 µL, 1.16 mmol), followed by $NEt_3$ (241 µL, 1.73 mmol) and the resultant mixture was stirred at RT for 18 h. It was then concentrated *in vacuo* and purification by column chromatography (gradient elution from $CH_2Cl_2$ to 10% MeOH in $CH_2Cl_2$) afforded the target product as a clear oil (119 mg, 0.452 mmol, 94%).

$^1$H NMR ($CDCl_3$, 600 MHz) $\delta_H$ 6.81-6.65 (m, 2H), 6.58-6.47 (m, 1H), 5.42-5.17 (m, 1H), 3.80 (d, 3H, $J$ = 16.5 Hz), 3.77-3.73 (m, 1H), 3.72 (d, 3H, $J$ = 3.8 Hz), 3.68-3.54 (m, 1H), 2.56-2.40 (m, 1H), 2.22-2.12 (m, 1H), 1.99-1.74 (m, 4H), 1.15 (t, 1H, $J$ = 7.5 Hz), 1.00 (t, 2H, $J$ = 7.5 Hz); $^{13}$C NMR ($CDCl_3$, 150 MHz) $\delta_C$ 173.3, 172.1, 153.7, 153.4, 150.5, 150.1, 132.8, 132.6, 112.9, 112.7, 111.8, 111.4, 111.2, 110.5, 56.5, 56.0, 55.9, 55.7, 55.7, 55.5, 47.6, 46.7, 34.1, 32.2, 28.0, 27.4, 23.6, 21.7, 9.2, 9.0; HRMS (ES+) calcd for $C_{15}H_{21}NO_3Na$ $[M+Na]^+$ 286.1419, observed 286.1408.

**3-(1-(2-chloroacetyl)pyrrolidin-2-yl)-4-methoxybenzoic acid (1)**

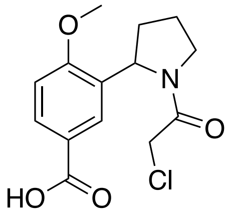

(S)-4-Methoxy-3-(pyrrolidin-2-yl)benzoic acid hydrochloride (100 mg, 0.388 mmol) was dissolved in DMF (8 mL). To this was added chloroacetyl chloride (37.0 µL, 0.465 mmol) at 0 °C, followed by $NEt_3$ (130 µL, 0.933 mmol) and the resultant mixture was stirred at RT for 18 h. It was then concentrated *in vacuo* and purification by column chromatography (gradient elution from $CH_2Cl_2$ with 1% AcOH to 10% MeOH in $CH_2Cl_2$ with 1% AcOH) afforded the target product as an off white solid (112 mg, 0.377 mmol, 97%).

$^1$H NMR ($CDCl_3$, 600 MHz) $\delta_H$ 8.09-7.98 (m, 1H), 7.76-7.69 (m, 1H), 6.99-6.91 (m, 1H), 5.47-5.35 (m, 1H), 4.18-4.11 (m, 1H), 4.00-3.84 (m, 4H), 3.84-3.70 (m, 2H), 2.41-2.25 (m, 1H), 2.03-1.84 (m, 3H); $^{13}$C NMR ($CDCl_3$, 150 MHz) $\delta_C$ 171.2, 170.6, 165.8, 165.4, 160.5, 160.1, 131.9, 131.2, 130.4, 130.2, 127.8, 127.3, 112.3, 121.4, 110.4, 110.1, 56.7, 55.6, 55.9, 55.7, 47.8, 47.8, 42.3, 41.8, 34.2, 32.0, 23.8, 21.7.

**(*S*)-3-(1-(2-chloroacetyl)pyrrolidin-2-yl)-4-methoxy-*N*-(pent-4-yn-1-yl)benzamide (NF686)**

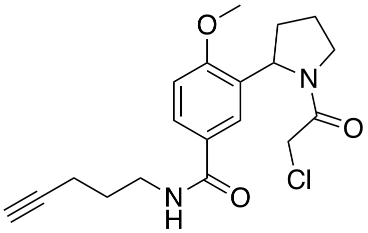

To **1** (50.0 mg, 0.168 mmol) in $CH_2Cl_2$ (5 mL) was added T3P (109 µL, 0.184 mmol, 50% solution in EtOAc), followed by $NEt_3$ (86.0 µL, 0.617 mmol) and 4-pentyn-1-amine (16.0 µL, 0.153 mmol). The resultant mixture was stirred at RT for 18 h, It was then concentrated *in vacuo* and purification by column chromatograhy (gradient elution from $CH_2Cl_2$ to 10% MeOH in $CH_2Cl_2$) afforded the target product as a clear oil (51.0 mg, 0.141 mmol, 92%).

$^1$H NMR ($CDCl_3$, 600 MHz) $\delta_H$ 7.87-7.65 (m, 1H), 7.51-7.39 (m, 1H), 6.98-6.87 (m, 1.5H), 6.44 (br s, 0.5H), 5.42-5.33 (m, 1H), 4.17-4.08 (m, 1H), 3.97-3.86 (m, 4H), 3.85-3.67 (m, 2H), 3.55-3.52 (m, 2H), 2.41-2.24 (m, 3H),

2.08-2.02 (m, 1H), 2.01-1.96 (m, 1H), 1.94-1.81 (m, 4H); $^{13}$C NMR ($CDCl_3$, 150 MHz) $\delta_C$ 167.5, 166.7, 165.7, 164.9, 158.7, 158.2, 130.0, 129.0, 127.4, 126.6, 124.1, 124.0, 110.5, 110.3, 84.0, 83.7, 69.3, 69.2, 56.7, 56.6, 55.8, 55.6, 47.9, 47.8, 42.5, 42.2, 39.4, 39.2, 34.4, 32.0, 28.1, 28.1, 23.9, 21.8, 16.4, 16.3; HRMS (ES+) calcd for $C_{19}H_{23}ClN_2O_3Na$ $[M+Na]^+$ 385.1295, observed 385.1286.

**_tert_-butyl 2-(2-methoxy-5-(trifluoromethyl)phenyl)-1_H_-pyrrole-1-carboxylate (2)**

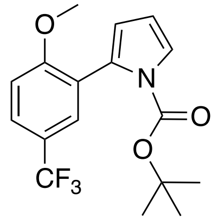

A mixture of 3-bromo-4-methoxybenzotrifluoride (1.02 g, 4.00 mmol), 1-Boc-pyrrole-2-boronic acid pinacol ester (1.76 g, 6.00 mmol) and $Cs_2CO_3$ (3.90 g, 12.0 mmol) in dioxane (16 mL)/ $H_2O$ (4 mL) was degassed for 20 min. XPhos Pd G4 (516 mg, 0.60 mmol) was then added and the mixture was further degassed for 10 min before it was heated to 100 °C for 16 h. Subsequent concentration *in vacuo* and purification by column chromatography (gradient elution from hexanes to 10% diethyl ether in hexanes) afforded the target product as a pale yellow oil (947 mg, 2.78 mmol, 70 %).

$^1$H NMR ($CDCl_3$, 600 MHz) $\delta_H$ 7.65-7.63 (m, 1H), 7.60 (d, 1H, $J$ = 2.3 Hz), 7.44 (dd, 1H, $J$ = 3.3, 1.8 Hz), 6.98 (d, 1H, $J$ = 8.6 Hz), 6.32 (t, 1H, $J$ = 3.3 Hz), 6.25 (dd, 1H, $J$ = 3.3, 1.8 Hz), 3.86 (s, 3H), 1.41 (s, 9H); $^{13}$C NMR ($CDCl_3$, 150 MHz) $\delta_C$ 159.9, 149.2, 129.6, 127.4 (q, $J$ = 3.6 Hz), 126.3 (q, $J$ = 3.9 Hz), 125.4, 124.9, 123.6, 122.7, 122.4, 122.4, 114.6, 110.5, 109.8, 83.2, 55.6, 27.5; HRMS (ES+) calcd for $C_{17}H_{18}F_3NO_3Na$ $[M+Na]^+$ 364.1136, observed 364.1125.

**2-(2-methoxy-5-(trifluoromethyl)phenyl)pyrrolidine (3)**

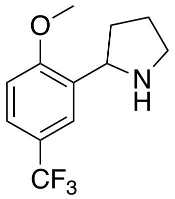

To **2** (100 mg, 0.293 mmol) in MeOH (10 mL) was added $Rh/Al_2O_3$ (30.0 mg, 5 wt.%). The resultant mixture was stirred under a $H_2$ atmosphere for 5 h, after which time it was filtered through a pad of celite and concentrated *in vacuo*. Conversion to the target pyrrolidine was confirmed by HRMS (ES+, calcd for $C_{17}H_{22}F_3NO_3Na$ $[2M+Na]^+$ 713.3003, observed 713.2988). The resultant product (101 mg, 0.293 mmol) was then dissolved in 4M HCl in dioxane (6 mL) and stirred at RT for 2 h. It was then concentrated *in vacuo* and used in the next steps without any further purification.

**2-chloro-1-(2-(2-methoxy-5-(trifluoromethyl)phenyl)pyrrolidin-1-yl)ethan-1-one (NF740)**

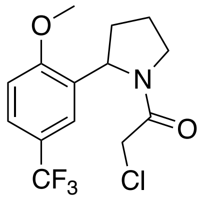

Compound **3** (105 mg, 0.304 mmol) was dissolved in DMF (5 mL). To this was added chloroacetyl chloride (36.0 µL, 0.453 mmol) at 0 °C, followed by $NEt_3$ (127 µL, 0.911 mmol) and the resultant mixture was stirred at RT for 18 h. It was then concentrated *in vacuo* and purification by column chromatography (gradient elution from $CH_2Cl_2$ to 30% diethyl ether in $CH_2Cl_2$) afforded the target product as an off white crystalline solid (65.0 mg, 0.202 mmol, 69%).

$^1$H NMR ($CDCl_3$, 600 MHz) $\delta_H$ 7.58-7.46 (m, 1H), 7.23-7.20 (m, 1H), 7.00-6.92 (m, 1H), 5.45-5.36 (m, 1H), 4.14-4.06 (m, 1H), 3.96-3.82 (m, 4H), 3.82-3.67 (m, 2H), 2.43-2.24 (m, 1H), 2.04-1.78 (m, 3H); $^{13}$C NMR ($CDCl_3$, 150 MHz) $\delta_C$ 165.6, 165.0, 158.6, 58.3, 131.2, 130.9, 127.2, 127.2, 126.8, 126.4 (q, *J* = 3.9 Hz), 125.5 (q, *J* = 3.9 Hz), 125.4, 125.0, 123.6, 123.4, 123.2, 123.0, 122.7 (q, *J* = 3.6 Hz), 122.6, 122.4, 122.3 (q, *J* = 3.7 Hz), 122.1, 121.8, 1221.4, 110.6, 110.4, 56.6, 56.4, 55.9, 55.6, 47.8, 47.7, 42.1, 41.8, 34.1, 32.0, 23.9, 21.7; HRMS (ES+) calcd for $C_{14}H_{15}ClF_3NO_2Na$ [2M+Na]$^+$ 665.1384, observed 665.1375.

**(*E*)-1-(2-(2-methoxy-5-(trifluoromethyl)phenyl)pyrrolidin-1-yl)-4-phenylbut-2-ene-1,4-dione (NF765)**

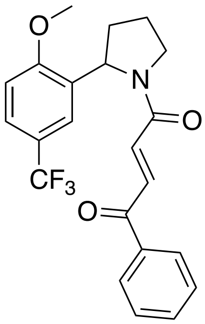

To trans-3-benzoylacrylic acid (42.0 mg, 0.238 mmol) in DMF (3 mL) was added HATU (130 mg, 0.342 mmol) and DIPEA (125 µL, 0.718 mmol), followed by **3** (45.0 mg, 0.184 mmol) in DMF (1 mL). The resultant mixture was stirred at RT for 18 h. It was then concentrated *in vacuo* and purified by column chromatography (gradient elution from hexanes to 30% diethyl ether in hexanes) to afford the target product as a yellow oil (38.0 mg, 0.094 mmol, 51%).

$^1$H NMR ($CDCl_3$, 600 MHz) $\delta_H$ 8.06-7.99 (m, 1H) 7.90-7.87 (m, 2H), 7.62-7.55 (m, 1H), 7.53-7.43 (m, 3H), 7.24-7.18 (m, 1H), 6.98-6.89 (m, 2H), 5.57-5.47 (m, 1H), 4.04-3.87 (m, 4H), 3.86-3.80 (m, 1H), 2.44-2.29 (m, 1H), 2.04-1.85 (m, 3H); $^{13}$C NMR ($CDCl_3$, 150 MHz) $\delta_C$ 189.9, 189.8, 164.0, 163.0, 158.6, 158.1, 137.0, 136.9, 136.9, 134.4, 134.0, 133.9, 133.7, 133.5, 133.1, 133.0, 131.9, 131.1, 128.9 (app q, *J* = 7.1 Hz), 128.8 (app q, *J* = 7.7 Hz), 128.7, 128.7, 126.3 (app q, *J* = 3.0), 125.6 (app q, *J* = 3.5 Hz), 123.2 (app q, *J* = 3.4 Hz), 123.1, 122.9, 122.7, 122.5 (app q, *J* = 4.2 Hz), 110.5, 110.5, 56.8, 56.4, 55.9, 55.7, 48.0, 47.5, 34.0, 32.1, 23.9, 21.8; HRMS (ES+) calcd for $C_{22}H_{20}F_3NO_3Na$ [M+Na]$^+$ 426.1293 observed 426.1283.

***tert*-butyl 2-(5-methoxy-2-(trifluoromethyl)phenyl)-1*H*-pyrrole-1-carboxylate (4)**

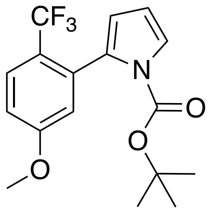

A mixture of 2-bromo-4-methoxy-1-(trifluoromethyl)benzene (692 mg, 2.71 mmol), 1-Boc-pyrrole-2-boronic acid pinacol ester (1.20 g, 4.09 mmol) and $Cs_2CO_3$ (2.60 g, 7.98 mmol) in dioxane (10.8 mL)/ $H_2O$ (2.7 mL) was degassed for 20 min. XPhos Pd G4 (348 mg, 0.40 mmol) was then added and the mixture was further degassed for 10 min before it was heated to 100 °C for 16 h. Subsequent concentration *in vacuo* and purification by column chromatography (gradient elution from hexanes to 10% diethyl ether in hexanes) afforded the target product as a pale yellow oil (723 mg, 2.12 mmol, 78%).

$^1$H NMR ($CDCl_3$, 600 MHz) $\delta_H$ 7.64 (d, 1H, *J* = 8.8 Hz), 7.44 (dd, 1H, *J* = 3.4, 1.8 Hz), 6.99 (ddd, 1H, *J* = 8.8, 2.7, 1.0 Hz), 6.92 (d, 1H, *J* = 2.6 Hz), 6.28 (t, 1H, *J* = 3.3 Hz), 6.25-6.14 (m, 1H), 3.87 (s, 3H), 1.29 (s, 9H); $^{13}$C NMR ($CDCl_3$, 150 MHz) $\delta_C$ 161.1, 148.9, 135.6 (q, *J* = 2.1 Hz), 129.4, 127.3 (q, *J* = 5.0 Hz), 125.2, 123.4, 122.5, 122.3, 122.0, 121.7, 118.0, 115.0, 113.1, 110.3, 83.3, 55.5, 27.4; HRMS (ES+) calcd for $C_{17}H_{18}F_3NO_3Na$ $[M+Na]^+$ 364.1136, observed 364.1124.

**2-(5-methoxy-2-(trifluoromethyl)phenyl)pyrrolidine (5)**

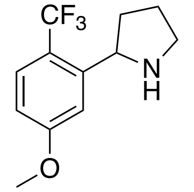

To **4** (100 mg, 0.293 mmol) in MeOH (10 mL) was added $Rh/Al_2O_3$ (30.0 mg, 5 wt.%). The resultant mixture was stirred under a $H_2$ atmosphere for 1.5 h, after which time it was filtered through a pad of celite and concentrated *in vacuo*. Conversion to the target pyrrolidine was confirmed by HRMS (ES+, calcd for $C_{17}H_{22}F_3NO_3Na$ $[M+Na]^+$ 368.1449, observed 368.1439). The resultant product (101 mg, 0.293 mmol) was then dissolved in 4M HCl in dioxane (6 mL) and stirred at RT for 2 h. It was then concentrated *in vacuo* and used in the next steps without any further purification.

**2-chloro-1-(2-(5-methoxy-2-(trifluoromethyl)phenyl)pyrrolidin-1-yl)ethan-1-one (NF741)**

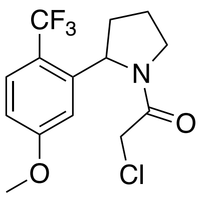

Compound **5** was dissolved in DMF (5 mL). To this was added chloroacetyl chloride (36.0 µL, 0.453 mmol) at 0 °C, followed by $NEt_3$ (127 µL, 0.911 mmol) and the resultant mixture was stirred at RT for 18 h. It was then concentrated *in vacuo* and purification by column chromatography (gradient elution from $CH_2Cl_2$ to 30% diethyl ether in $CH_2Cl_2$) afforded the target product as a white solid (53.0 mg, 0.165 mmol, 56%).

$^{1}H$ NMR ($CDCl_3$, 600 MHz) $\delta_H$ 7.66-7.58 (m, 1H), 6.89-6.72 (m, 2H), 5.46-5.34 (m, 1H), 4.14-4.03 (m, 1H), 4.00-3.87 (m, 1H), 3.88-3.59 (m, 5H), 2.56-2.36 (m, 1H), 2.16-1.79 (m, 3H); $^{13}C$ NMR ($CDCl_3$, 150 MHz) $\delta_C$ 165.6, 164.7, 162.6, 162.2, 143.9, 143.7, 128.7 (q, $J$ = 5.9 Hz), 128.3 (q, $J$ = 5.9 Hz), 127.4, 127.2, 125.6, 125.4, 123.8, 123.6, 122.0, 121.8, 119.6, 119.4, 119.2, 119.2, 119.0, 118.8, 118.6, 113.2, 111.7, 111.5, 110.9, 58.1 (q, $J$ = 2.4 Hz), 57.8 (q, $J$ = 2.2 Hz), 55.6, 55.4, 48.4, 48.3, 42.4, 42.0, 36.1, 34.0, 24.1, 21.5; HRMS (ES+) calcd for $C_{14}H_{15}ClF_3NO_2H$ $[M+H]^+$ 322.0822, observed 322.0814.

**(*E*)-1-(2-(5-methoxy-2-(trifluoromethyl)phenyl)pyrrolidin-1-yl)-4-phenylbut-2-ene-1,4-dione (NF764)**

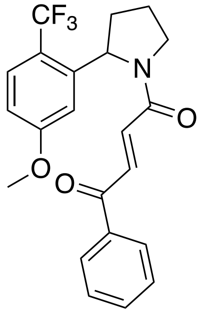

To trans-3-Benzoylacrylic acid (42.0 mg, 0.238 mmol) in DMF (3 mL) was added HATU (130 mg, 0.342 mmol) and DIPEA (125 µL, 0.718 mmol), followed by **5** (45.0 mg, 0.184 mmol) in DMF (1 mL). The resultant mixture was stirred at RT for 18 h. It was then concentrated *in vacuo* and purified by column chromatography (gradient elution from $CH_2Cl_2$ to 30% diethyl ether in $CH_2Cl_2$) to afford the target product as an orange oil (49.0 mg, 0.122 mmol, 66%).

$^{1}H$ NMR ($CDCl_3$, 600 MHz) $\delta_H$ 8.04-8.01 (m, 1H), 7.86-7.79 (m, 2H), 7.64-7.60 (m, 1H), 7.58-7.42 (m, 3H), 6.92-6.90 (m, 1H), 6.85-6.72 (m, 2H), 5.57-5.48 (m, 1H), 4.06-3.86 (m, 2H), 3.82-3.81 (m, 3H), 2.57-2.42 (m, 1H), 2.20-1.85 (m, 3H); $^{13}C$ NMR ($CDCl_3$, 150 MHz) $\delta_C$ 189.8, 189.6, 164.0, 162.9, 162.5, 162.2, 114.4, 144.2, 136.9, 136.9, 134.6, 134.4, 133.7, 133.4, 133.0, 132.7, 128.8, 128.8, 128.7, 128.7, 128.3, 125.5, 123.7, 119.6, 119.4, 118.9, 118.7, 113.7, 112.6, 111.3, 110.3, 58.3 (app q, $J$ = 2.4 Hz), 58.1 (q, $J$ = 2.3 Hz), 55.5, 55.3, 48.4, 48.1, 36.0, 34.1, 24.0, 21.7; HRMS (ES+) calcd for $C_{22}H_{20}F_3NO_3H$ $[M+H]^+$ 404.1474, observed 404.1466.

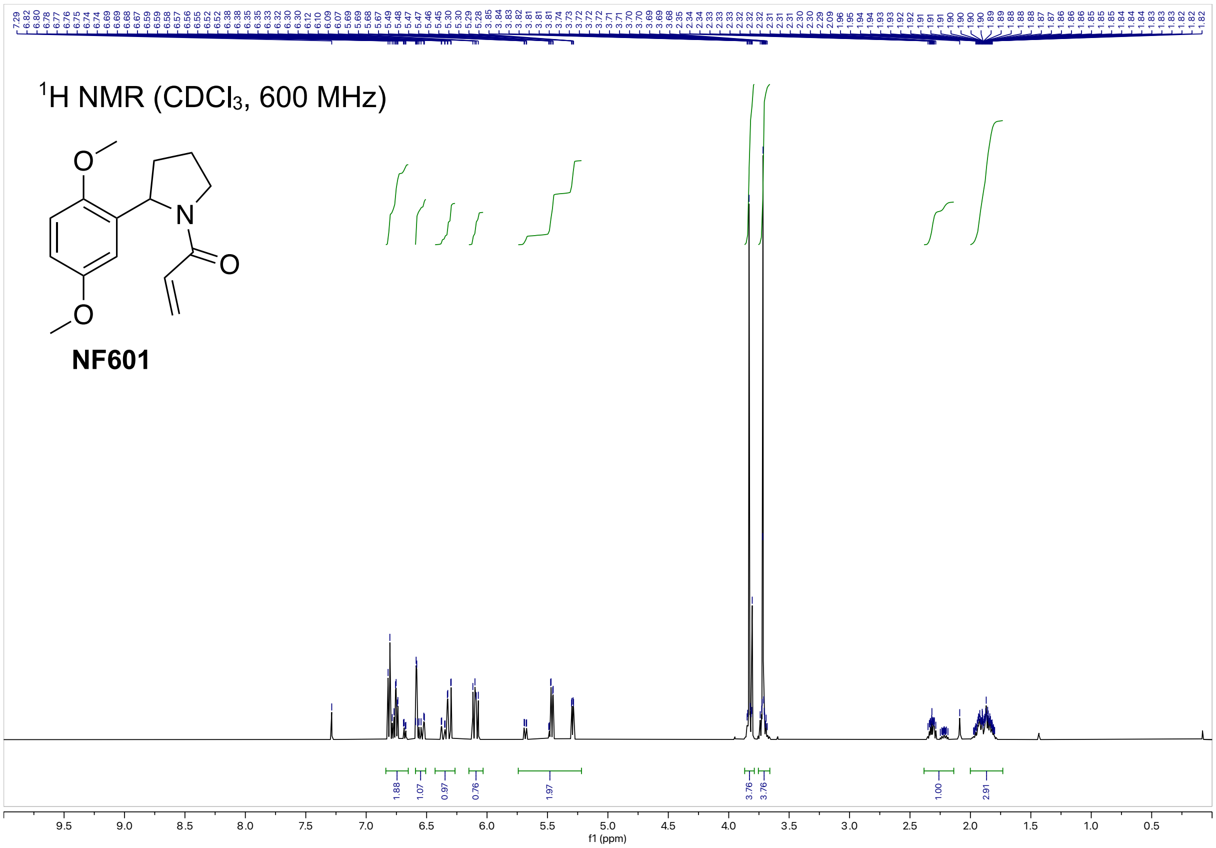

$^1$H NMR (CDCl$_3$, 600 MHz)
NF601
f1 (ppm)


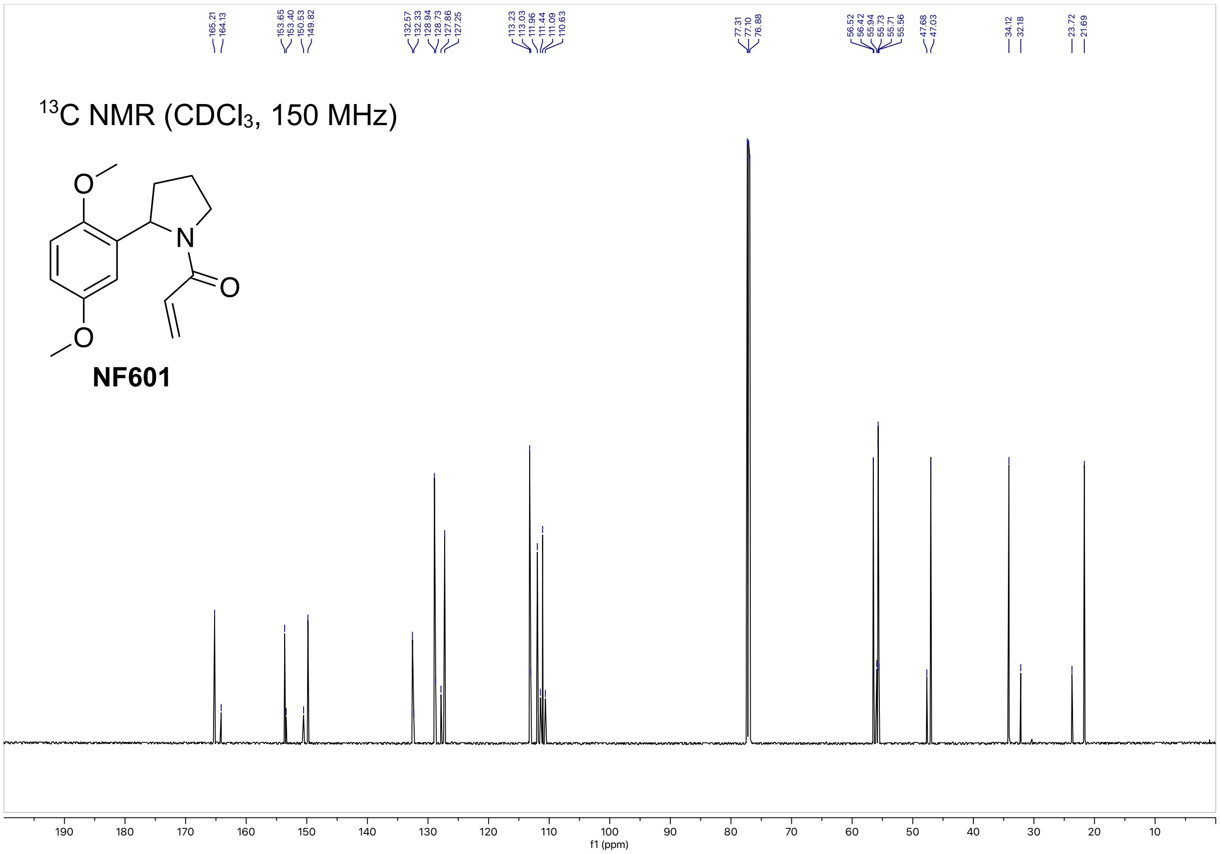

165.21
164.13
153.65
153.40
150.53
149.82
132.57
132.33
128.94
128.73
127.86
127.25
113.23
113.03
111.96
111.44
111.09
110.63
77.31
77.10
76.88
56.52
56.42
55.94
55.73
55.71
55.56
47.68
47.03
34.12
32.18
23.72
21.69
$^{13}$C NMR (CDCl$_3$, 150 MHz)
NF601
f1 (ppm)

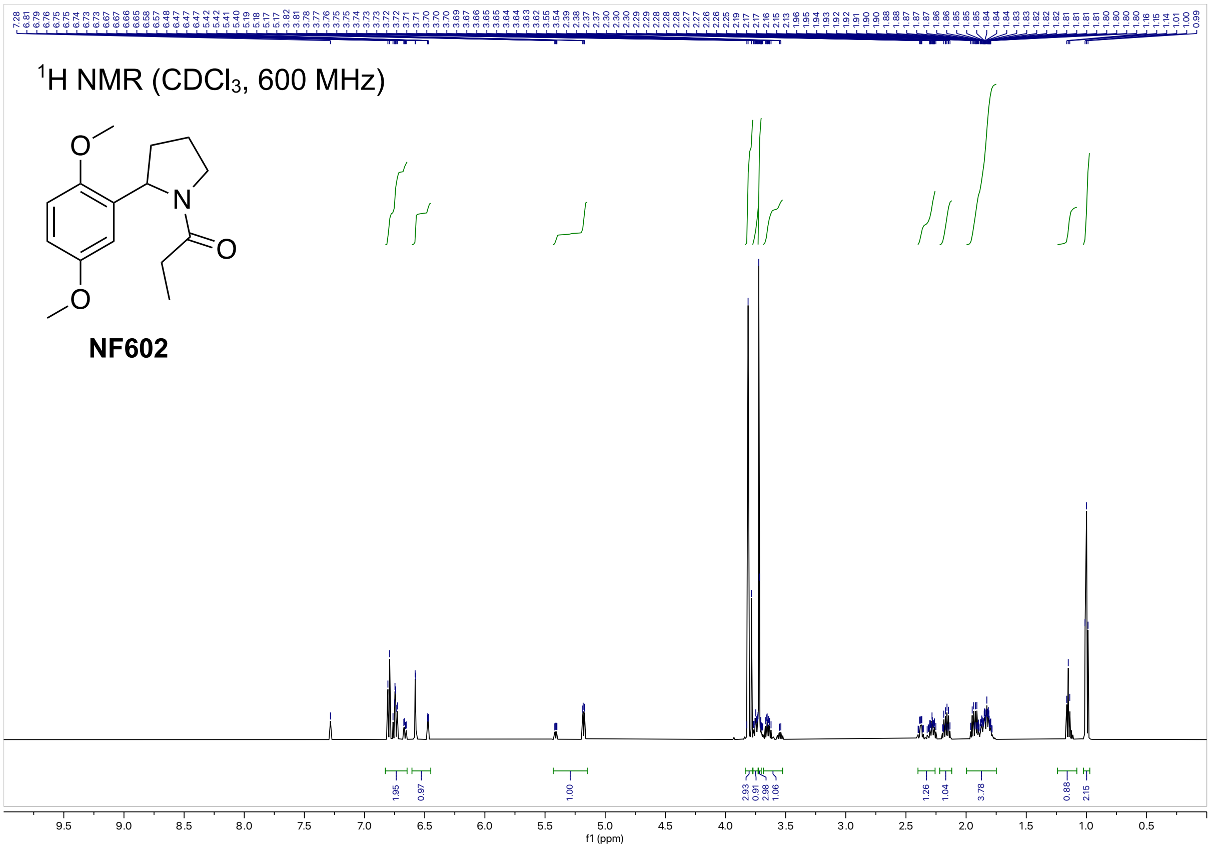

1H NMR (CDCl3, 600 MHz)
NF602
1.95
0.97
1.00
2.93
0.91
2.98
1.06
1.26
1.04
3.78
0.88
2.15
f1 (ppm)


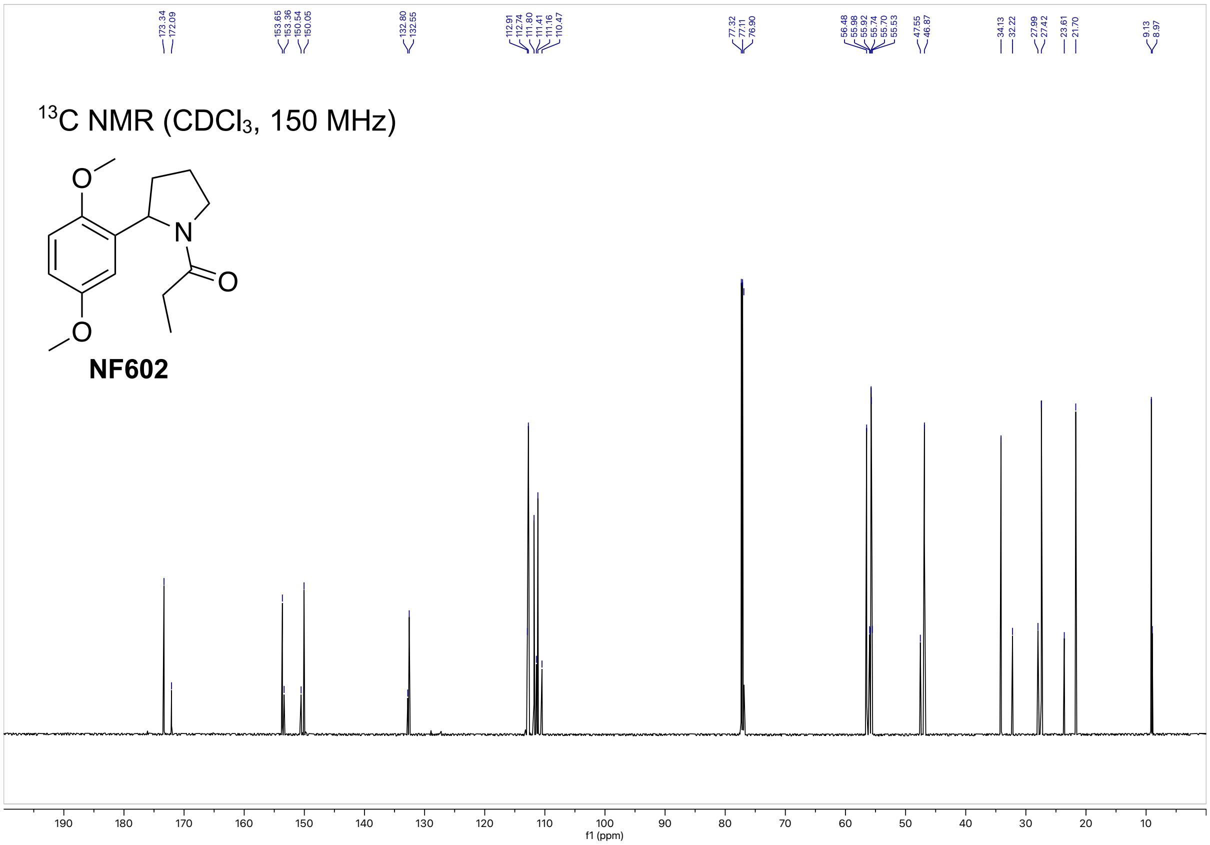

13C NMR (CDCl3, 150 MHz)
NF602
173.34
172.09
153.65
153.36
150.54
150.05
132.80
132.55
112.91
112.74
111.80
111.41
111.16
110.47
77.32
77.11
76.90
56.48
55.98
55.92
55.74
55.70
55.53
47.55
46.87
34.13
32.22
27.99
27.42
23.61
21.70
9.13
8.97
f1 (ppm)

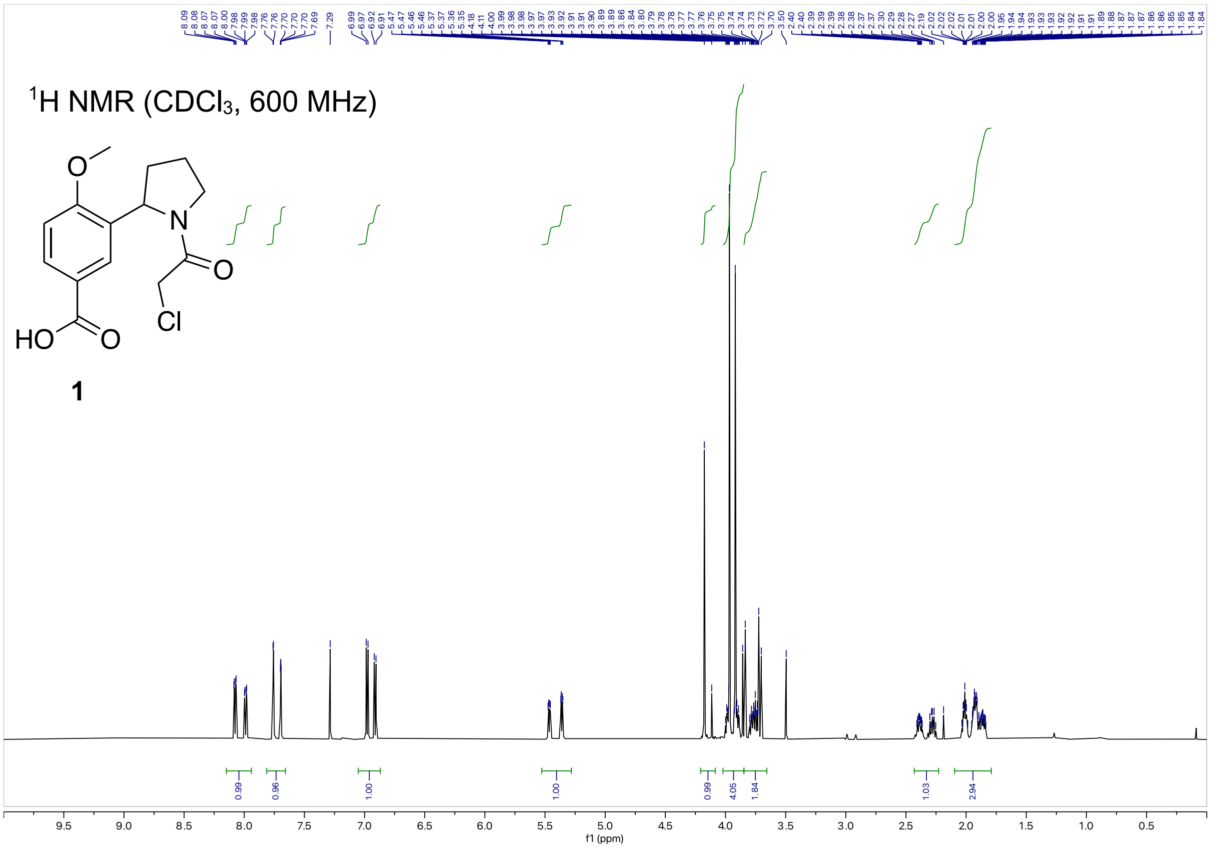

$^1$H NMR ($CDCl_3$, 600 MHz)
O
N
O
HO
O
Cl
1
f1 (ppm)


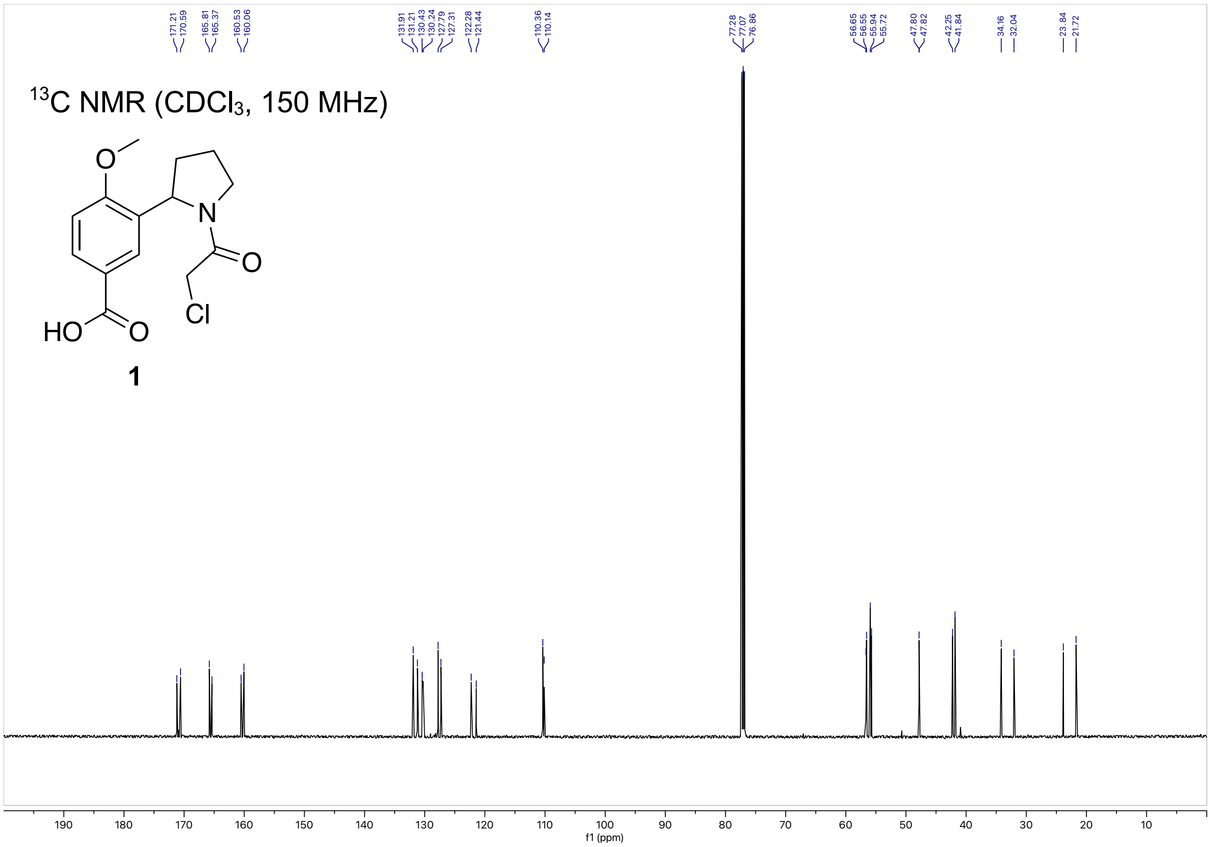

$^{13}$C NMR ($CDCl_3$, 150 MHz)
O
N
O
HO
O
Cl
1
f1 (ppm)

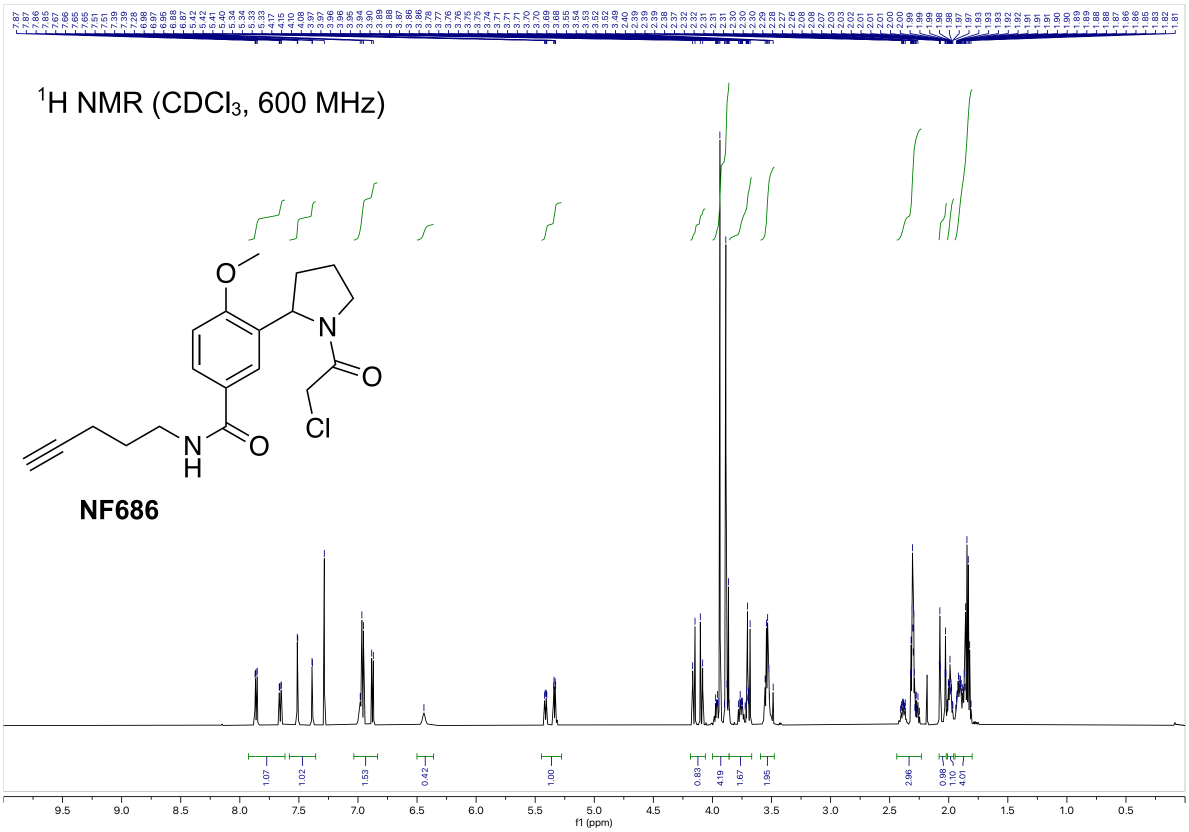

¹H NMR (CDCl₃, 600 MHz)
NF686
Cl
f1 (ppm)


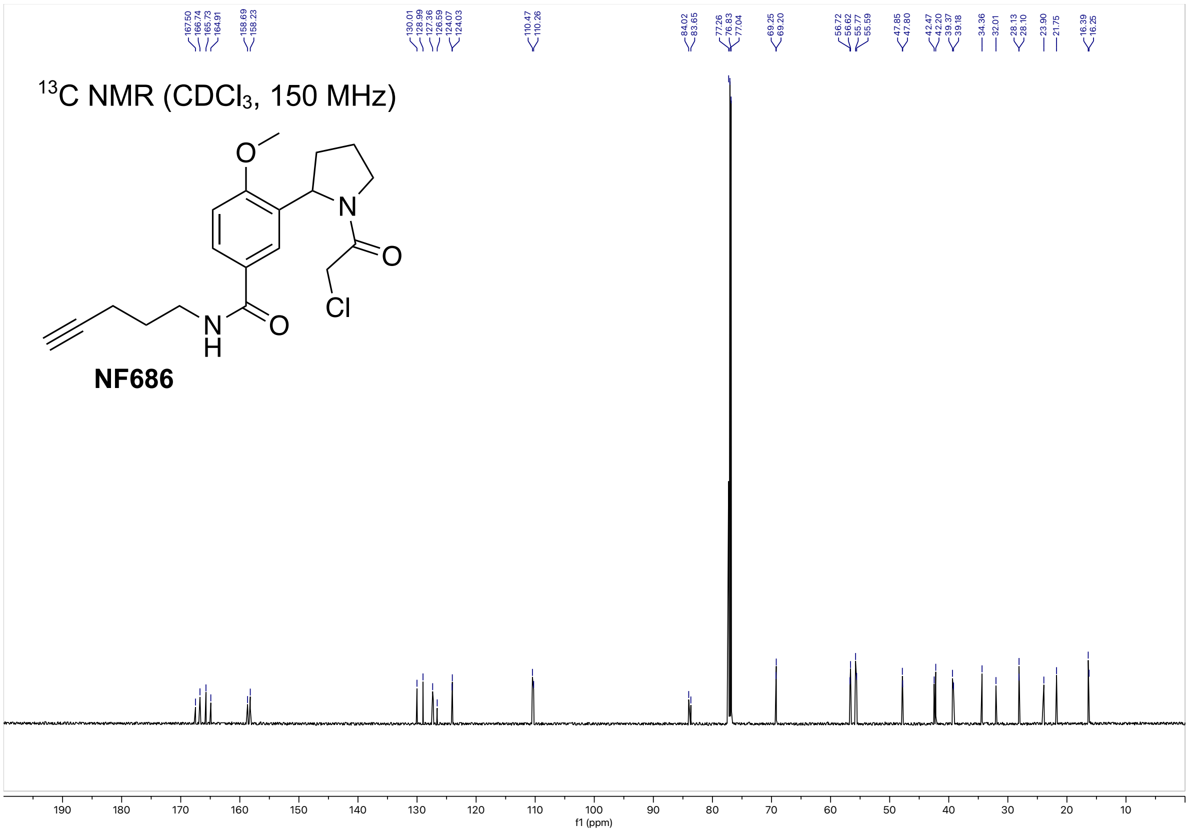

¹³C NMR (CDCl₃, 150 MHz)
NF686
Cl
f1 (ppm)

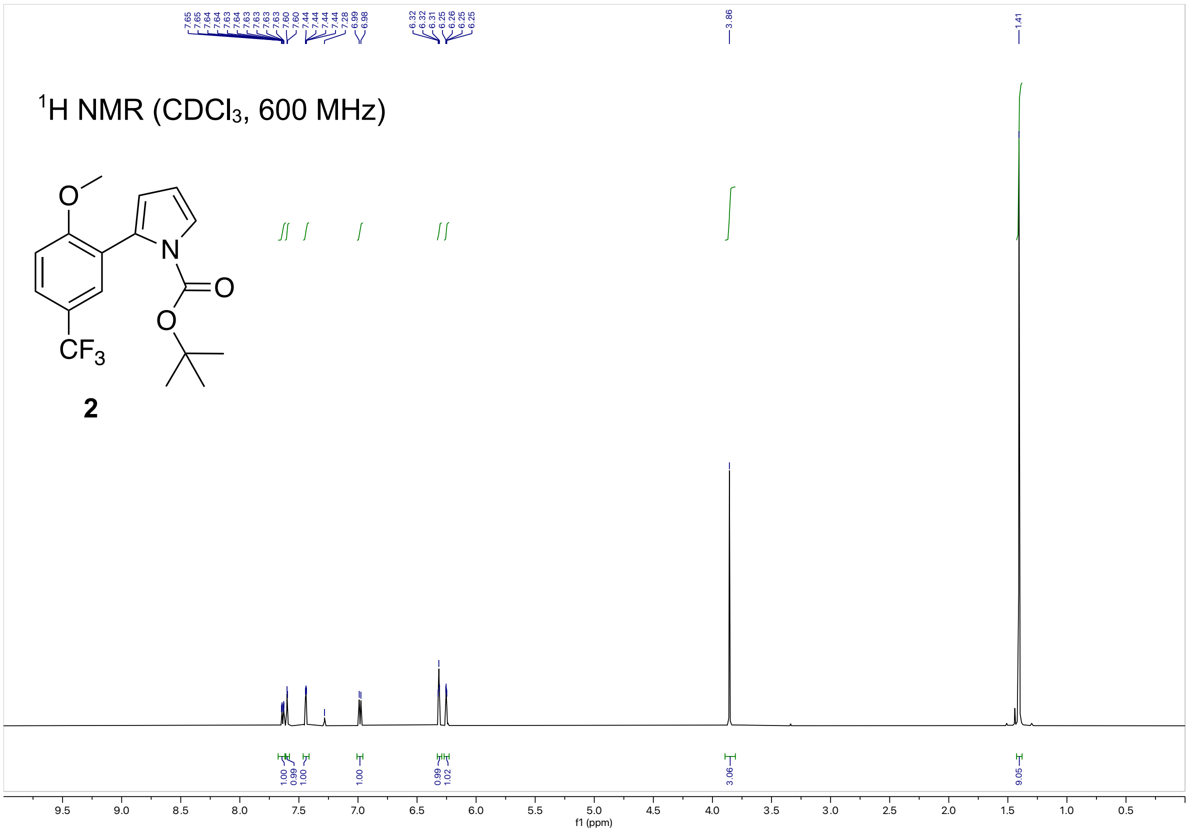

[1]H NMR (CDCl3, 600 MHz)
CF3
2
7.65 7.65 7.64 7.63 7.64 7.63 7.63 7.63 7.60 7.60 7.44 7.44 7.44 7.44 7.28 6.99 6.98
6.32 6.32 6.31 6.25 6.26 6.25 6.25
3.86
1.41
1.00 0.99 1.00 1.00 0.99 1.02 3.06 9.05
9.5 9.0 8.5 8.0 7.5 7.0 6.5 6.0 5.5 5.0 4.5 4.0 3.5 3.0 2.5 2.0 1.5 1.0 0.5
f1 (ppm)


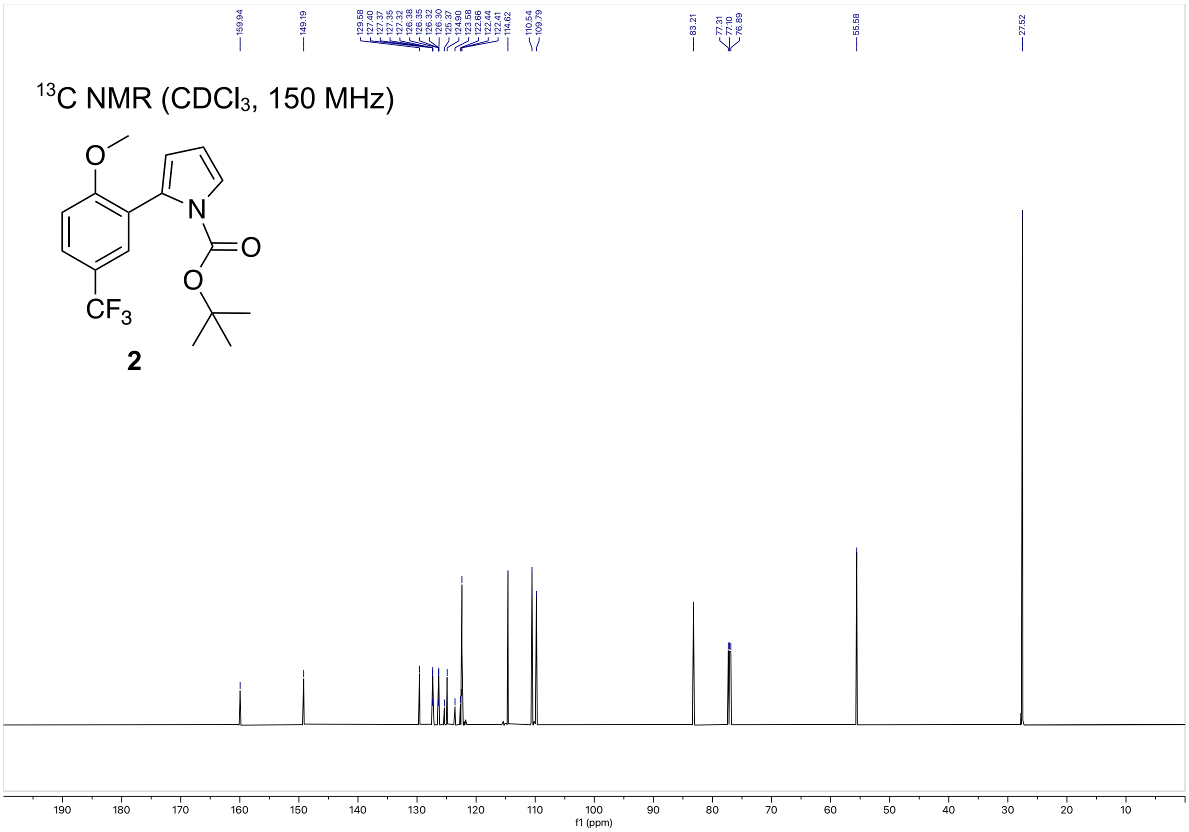

[13]C NMR (CDCl3, 150 MHz)
CF3
2
159.94
149.19
129.58 127.40 127.37 127.35 127.32 126.38 126.35 126.32 126.30 125.37 124.90 123.58 122.66 122.44 122.41 114.62
110.54 109.79
83.21
77.31 77.10 76.89
55.58
27.52
190 180 170 160 150 140 130 120 110 100 90 80 70 60 50 40 30 20 10
f1 (ppm)

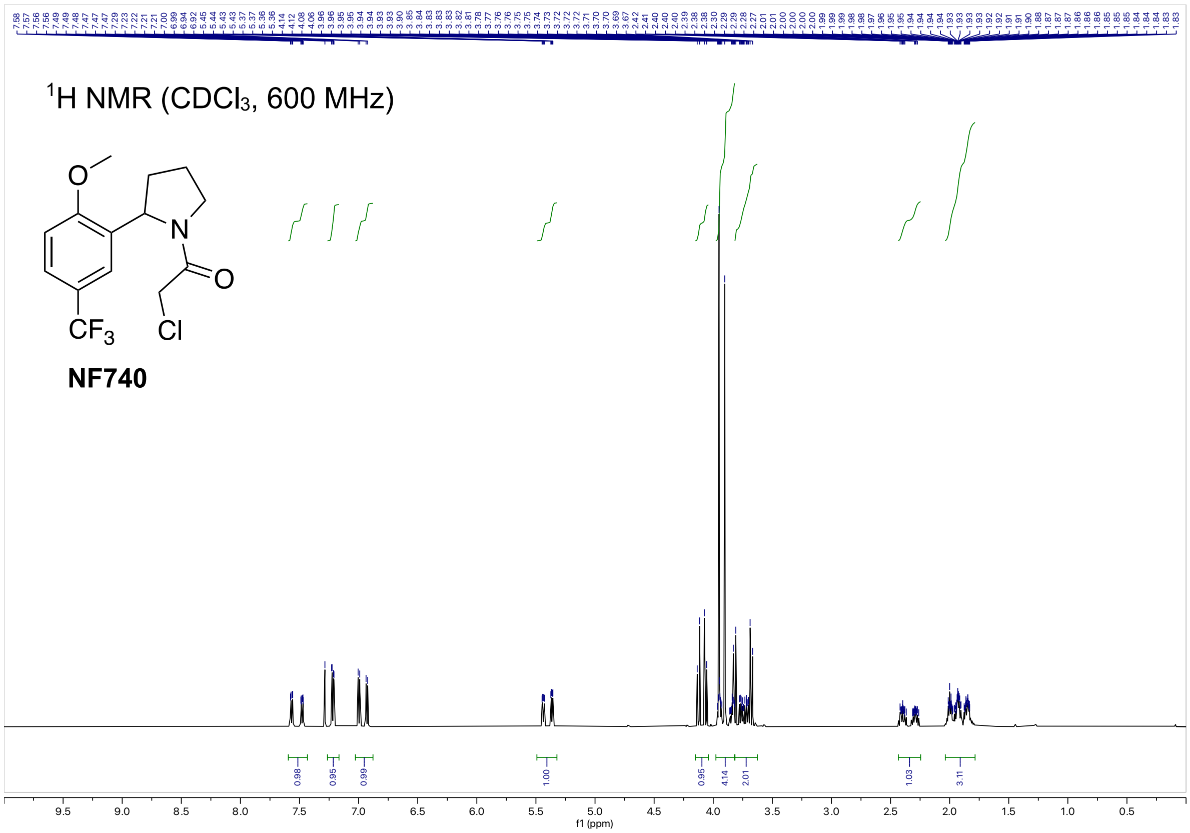

1H NMR (CDCl3, 600 MHz)
NF740
f1 (ppm)


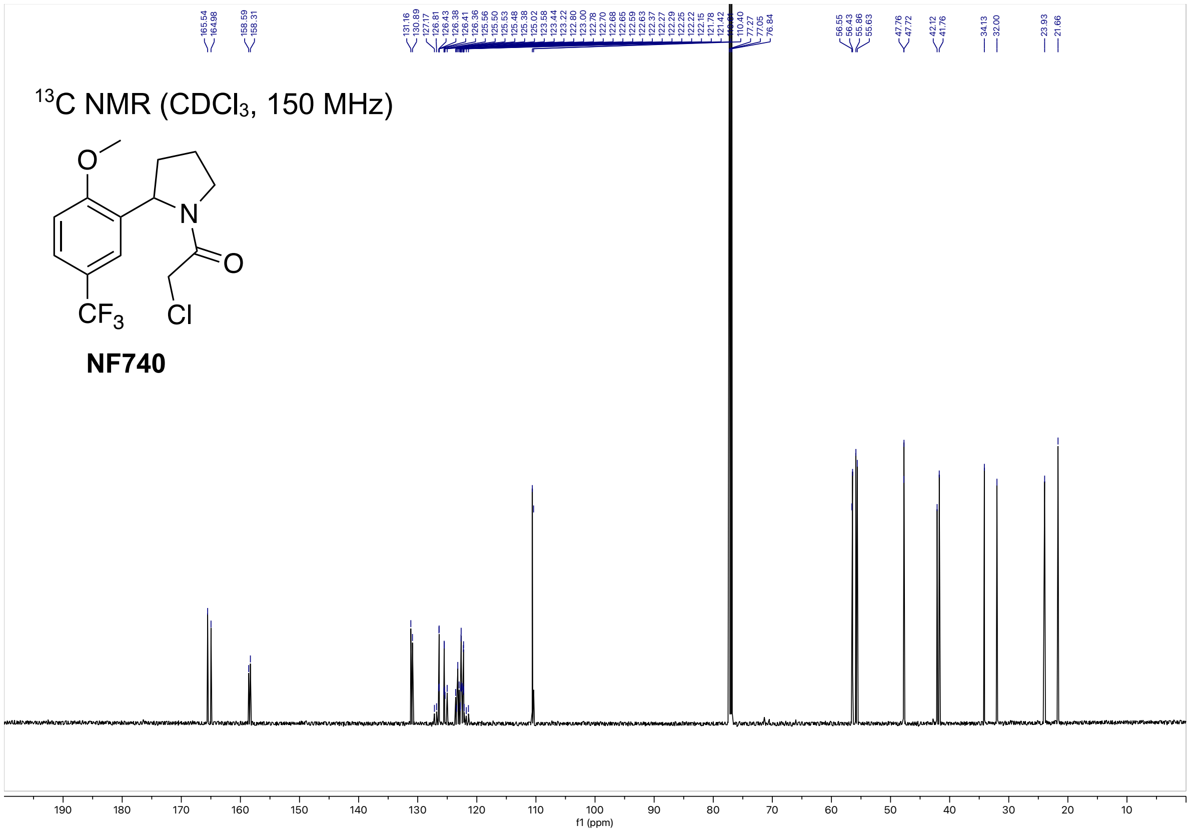

13C NMR (CDCl3, 150 MHz)
NF740
f1 (ppm)

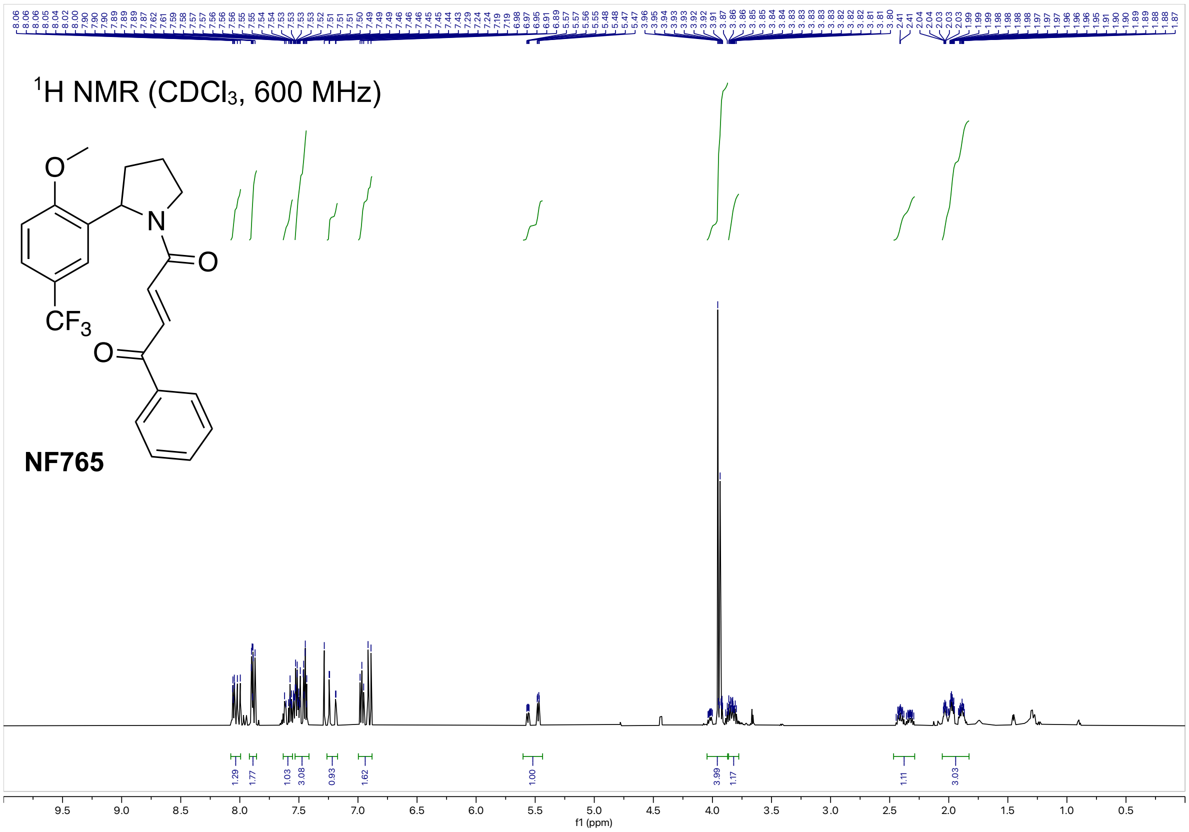

$^1$H NMR (CDCl$_3$, 600 MHz)
NF765
f1 (ppm)


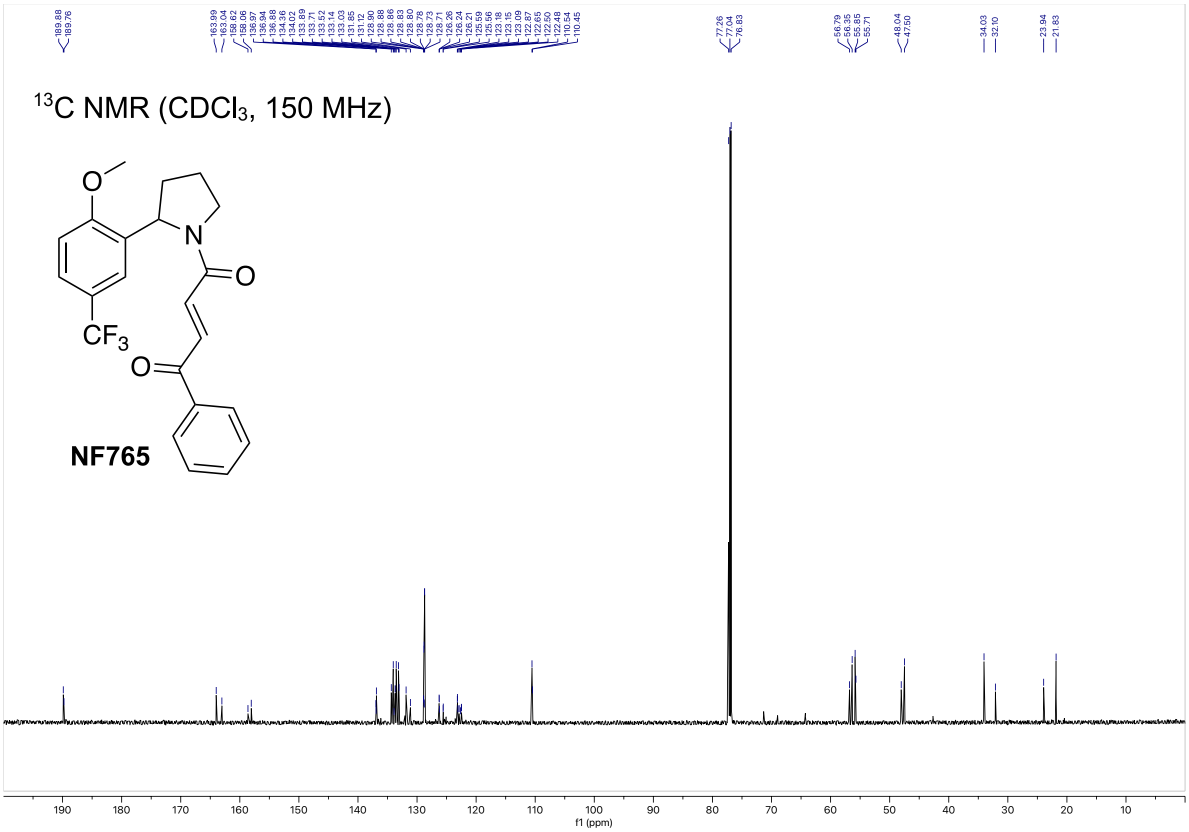

$^{13}$C NMR (CDCl$_3$, 150 MHz)
NF765
f1 (ppm)

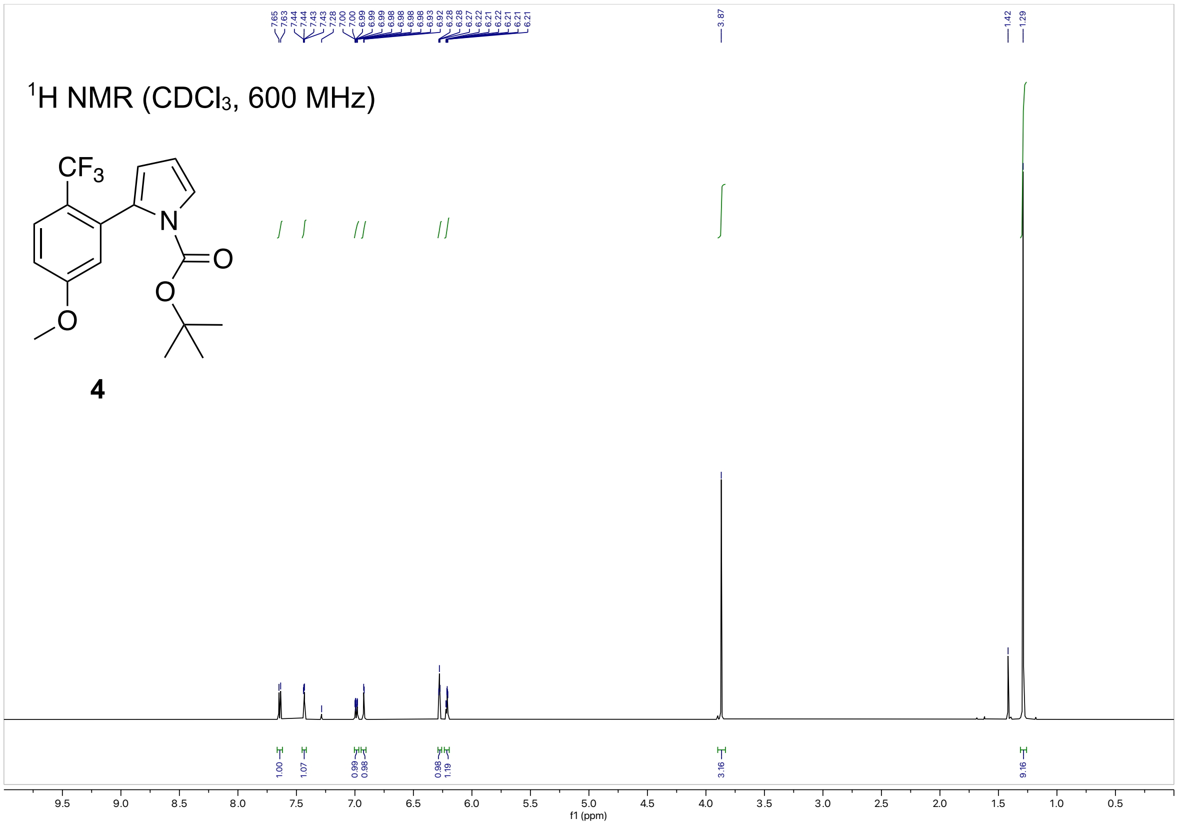
1H NMR (CDCl3, 600 MHz)
CF3
N
O
O
O
4
f1 (ppm)

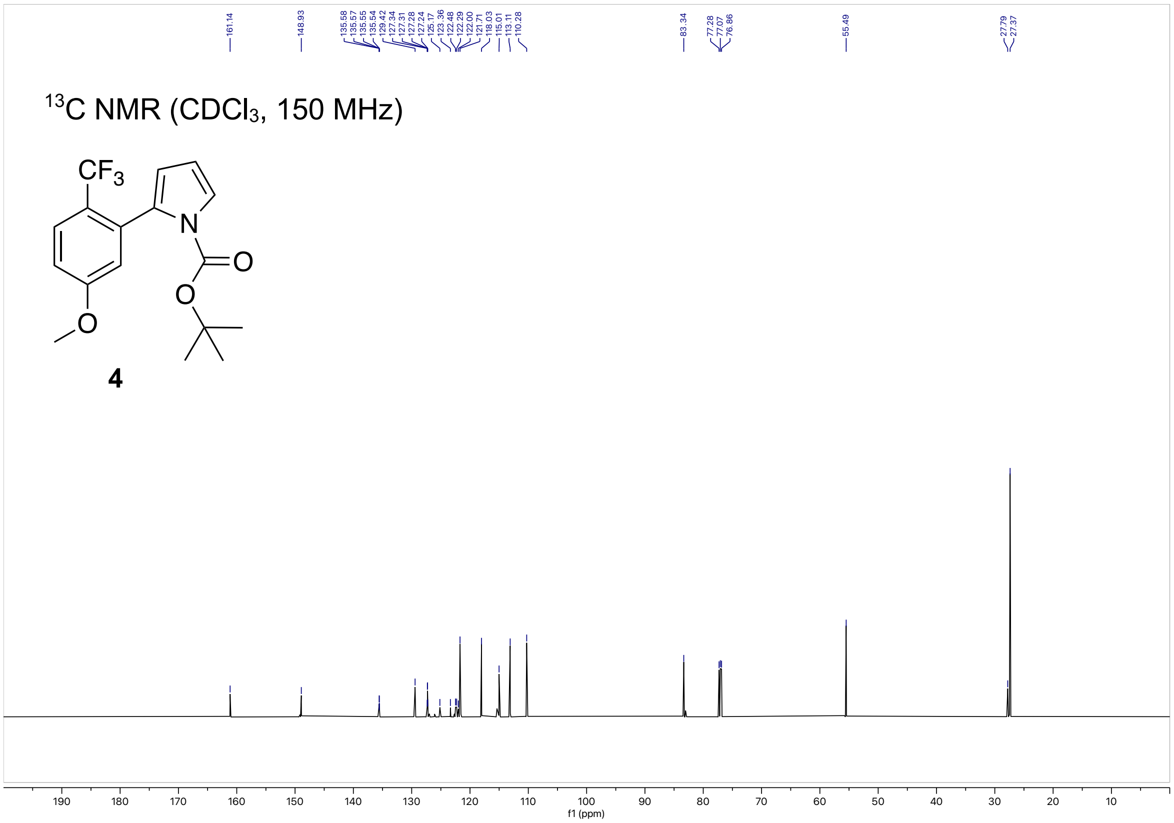
13C NMR (CDCl3, 150 MHz)
CF3
N
O
O
O
4
f1 (ppm)

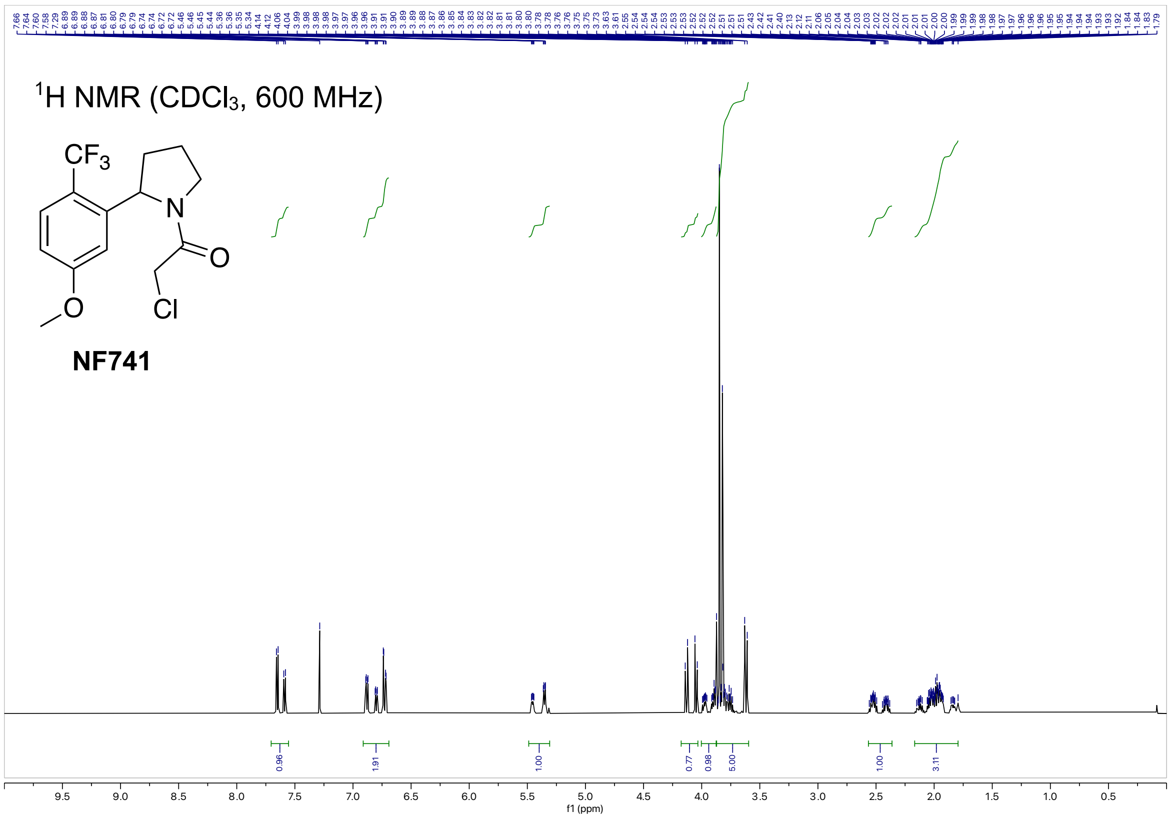

$^{1}$H NMR (CDCl$_3$, 600 MHz)
NF741
0.96
1.91
1.00
0.77
0.98
5.00
1.00
3.11
f1 (ppm)


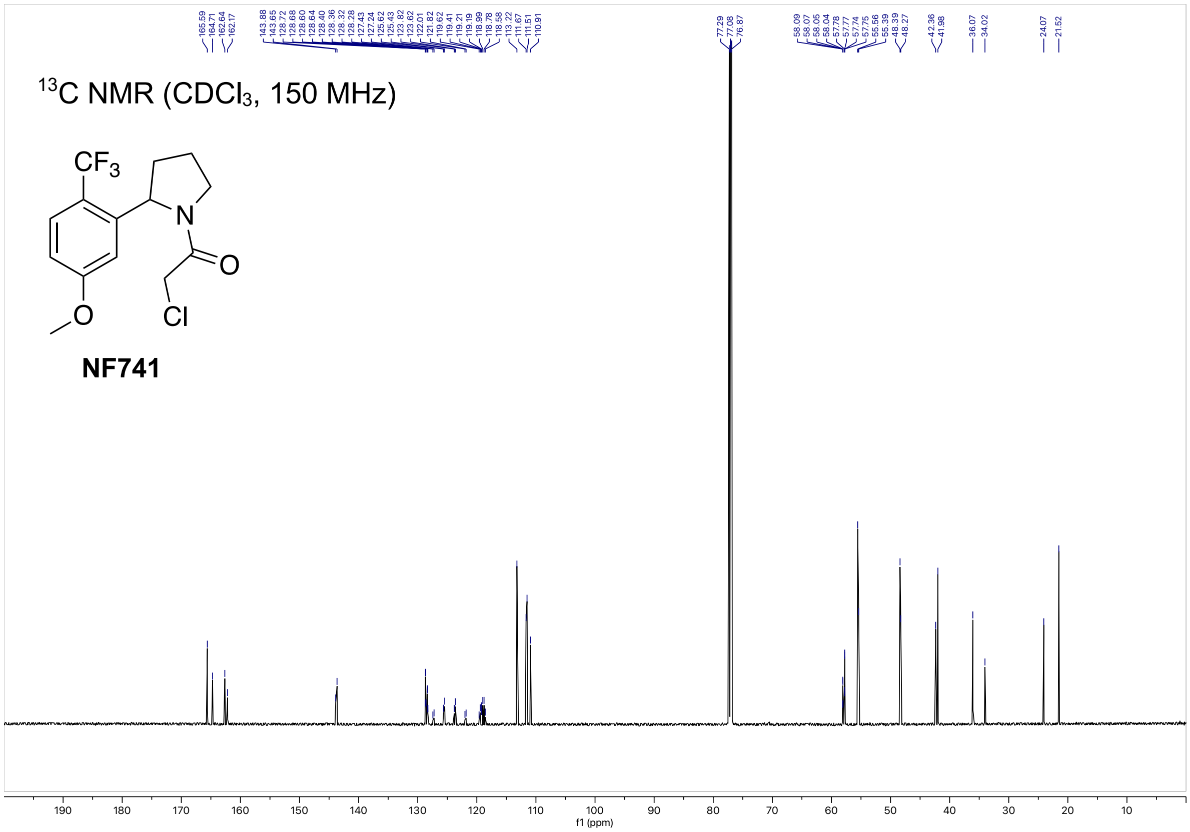

$^{13}$C NMR (CDCl$_3$, 150 MHz)
NF741
f1 (ppm)

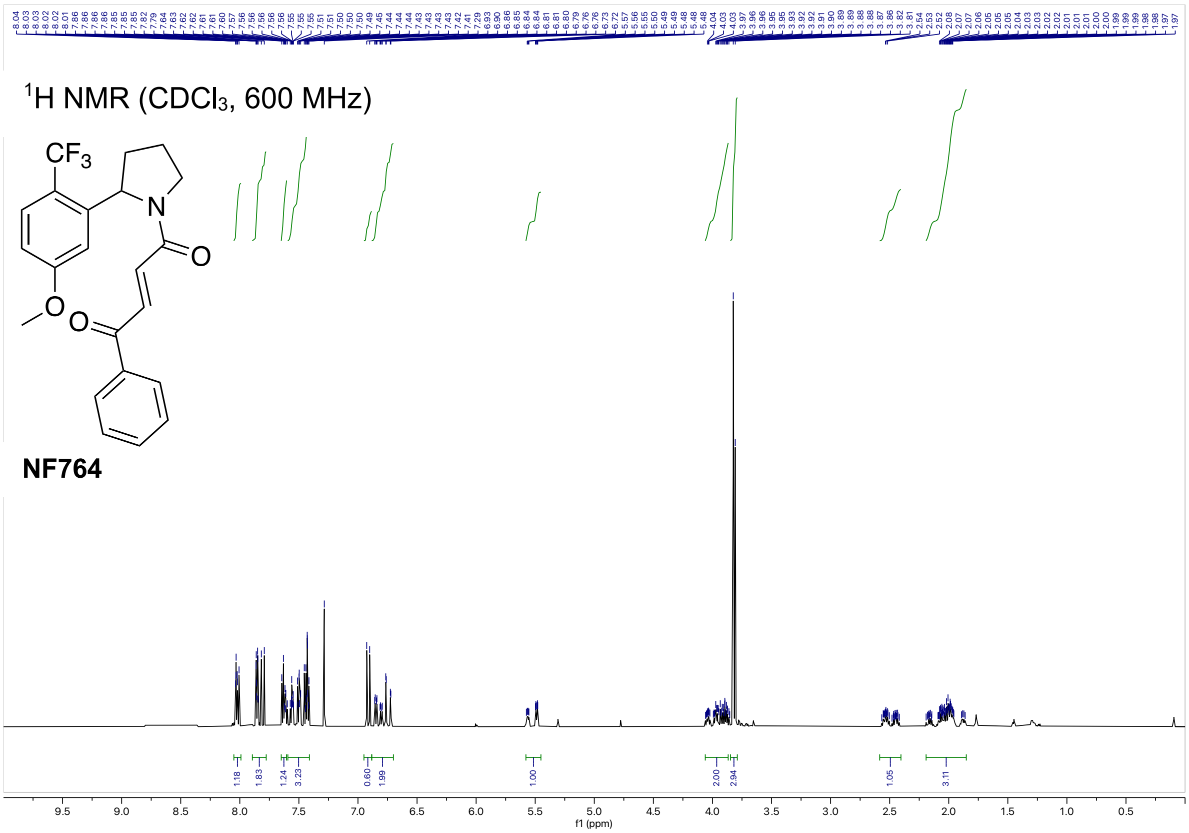

$^1$H NMR ($CDCl_3$, 600 MHz)
CF3
N
O
O
NF764
f1 (ppm)


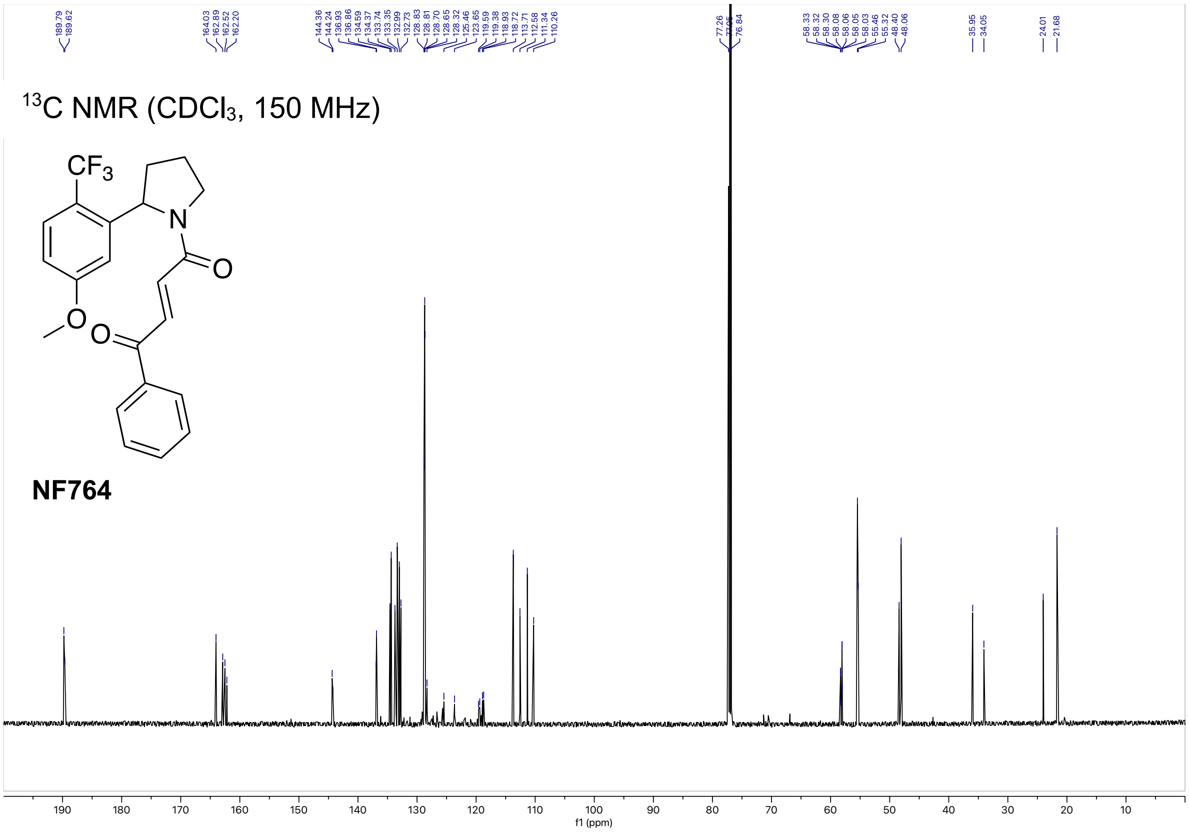

$^{13}$C NMR ($CDCl_3$, 150 MHz)
CF3
N
O
O
NF764
f1 (ppm)
